# Unraveling the Molecular Mechanisms of ABHD5 Membrane Targeting

**DOI:** 10.1101/2025.05.06.652485

**Authors:** Amit Kumar, Matthew Sanders, Huamei Zhang, Li Zhou, Shahnaz Parveen, Miriam C. Jensen, Thomas J.D. Jørgensen, Christopher V. Kelly, James G. Granneman, Yu-ming M. Huang

**Affiliations:** Department of Physics and Astronomy, Wayne State University, Detroit, MI 48201, USA; Center for Molecular Medicine and Genetics, School of Medicine, Wayne State University, Detroit, MI 48201, USA; Center for Integrative Metabolic and Endocrine Research, School of Medicine, Wayne State University, Detroit, MI 48201, USA; Department of Biochemistry and Molecular Biology, Southern Denmark University, Odense Denmark; Barber Center for Multiscale Systems Biology, Detroit, MI 48202, USA

**Keywords:** LD-associated protein activation, lipolysis regulation, lipid homeostasis, lipid droplet biophysics, protein-membrane interaction, Gaussian accelerated molecular dynamics

## Abstract

ABHD5 is a master regulator of PNPLA family lipases, particularly PNPLA2 (ATGL), the rate-limiting triglyceride (TAG) hydrolase in metabolic tissues. Despite its central role in lipid metabolism, the molecular basis by which ABHD5 recognizes membranes and regulates enzymatic activity remains poorly understood. Here, we report an integrated computational-experimental study revealing how α/β-hydrolase domain-containing protein 5 (ABHD5) dynamically engages lipid droplet (LD) and endoplasmic reticulum membranes. Using multiscale molecular dynamics simulations, hydrogen-deuterium exchange mass spectrometry, and site-directed mutagenesis, we uncover a sequential dual-site membrane recognition mechanism, where the N-terminus provides initial anchoring and a lid helix within the insertion segment forms a crucial secondary contact. Membrane binding triggers a dramatic conformational switch in this lid, expanding the pseudosubstrate pocket and stabilizing an open conformation of ABHD5. This structural transition is coupled to membrane remodeling, inducing localized curvature and forming a triacylglycerol-enriched nanodomain beneath the ABHD5 pseudosubstrate pocket. This bidirectional interaction between ABHD5 and the membrane provides molecular insights into the structural basis of ABHD5 membrane association and lipid remodeling. Our findings establish new principles for how LD-binding proteins achieve functional specificity through membrane-dependent regulation, offering novel molecular targets for interventions in metabolic diseases.

**Teaser:** A key protein controls fat breakdown by changing shape when it binds membranes and reshapes local fat droplets.

## Introduction

Lipid metabolism dysregulation underlies major global health challenges, from the obesity epidemic affecting over 650 million adults worldwide to cardiovascular diseases that claim 20 million lives annually (1–6). At the cellular level, lipids function as essential membrane components, signaling molecules, and energy reservoirs, with their precise regulation determining cellular fate and organismal health (7–10). Despite this fundamental importance, our mechanistic understanding of how lipid homeostasis is dynamically controlled across different cellular compartments remains incomplete, limiting therapeutic interventions for metabolic diseases.

α/β-Hydrolase domain-containing protein 5 (ABHD5) has emerged as a master regulator orchestrating diverse lipid metabolic pathways with remarkable tissue specificity. It activates triglyceride (TAG) lipolysis in adipocytes and cardiomyocytes (11, 12), drives skin barrier formation in keratinocytes (13, 14), and regulates phospholipid remodeling in hepatocytes (15). Critically, loss of ABHD5 function in humans causes Chanarin-Dorfman syndrome, whose diverse pathological features include severe ichthyosis, hepatomegaly, and progressive myopathy (13).

ABHD5 is a pseudoenzyme that evolved from the duplication of ABHD4, a phospholipase, and the inactivation of the catalytic serine. Thus, although ABHD5 is an essential activator of certain patatin-like phospholipase domain-containing (PNPLA) family members, ABHD5 itself lacks intrinsic catalytic activity (16, 17). At a basic level, ABHD5 enables access for diverse PNPLA paralogs to their specific membrane-delimited substrates; however, the mechanisms involved are not understood (13, 18, 19). ABHD5 selectively binds to intracellular lipid droplets (LD), specialized cellular organelles composed of a phospholipid monolayer surrounding a triacylglycerol-rich neutral lipid core. We hypothesize that ABHD5 binding alters the nanoscale chemical and biophysical properties of the LD monolayer through direct lipid interactions and membrane remodeling. Understanding how ABHD5 engages and remodels LD surfaces is essential for elucidating the molecular basis of its membrane-dependent regulatory functions.

ABHD5 must bind membranes to activate PNPLA2 (20); however, it is presently unknown how ABHD5 targets and orients on membranes, limiting our understanding of its function at the membrane interface, where its biological activity occurs. Prior structural investigations have provided only partial insights. NMR studies characterized the intrinsically disordered N-terminal region (residues 1-43), which may contribute to membrane association (21, 22). Homology and deep learning-based models captured aspects of the soluble ABHD5 structure, including the canonical hydrolase fold and a unique insertion segment (residues 180-270) that contains a short lid helix (residues 198-207) near the pseudosubstrate pocket as part of the membrane insertion domain (residues 180-230) (23) (**Fig. 1**). However, these models are limited to solution-phase conformations and do not address how ABHD5 engages with lipid layers, leaving its membrane-bound configuration and mechanism of action unresolved.

**Figure 1:**
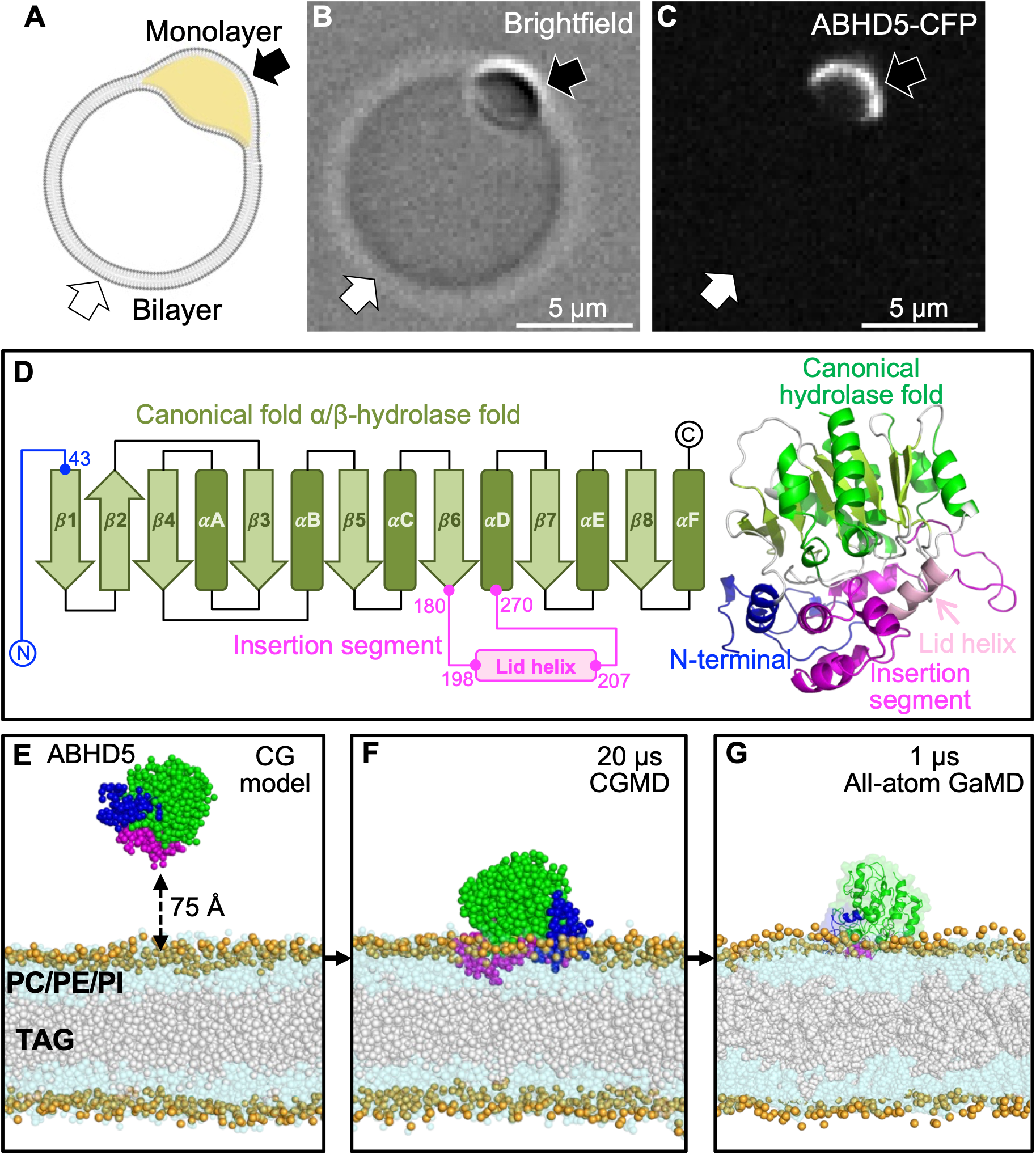
Multiscale simulations combined with experiments identify the protein motifs responsible for membrane binding. ABHD5 selectively binds to droplet-embedded vesicles (DEVs). Schematic of a DEV consisting of a phospholipid bilayer (gray) enclosing a LD monolayer (yellow) containing TAG. The white arrow indicates the bilayer, and the black arrow indicates the monolayer (A). Brightfield image of a DEV showing the phase-separated bilayer (white arrow) and droplet monolayer (black arrow) (B). Fluorescence image of ABHD5-CFP bound to the DEV, showing strong enrichment at the TAG-containing monolayer (black arrow) and absence from the bilayer (white arrow) (C). Our computational workflow incorporates validating and equilibrating the AlphaFold protein structure (D), converting this structure to a CG model (E), simulating the protein conformation changes and membrane remodeling with CGMD (F), and back-conversion and equilibration of the structures with GaMD (G).

This structural gap is particularly striking given that ABHD5 activates multiple PNPLA paralogs that localize to distinct cellular membranes. For example, it regulates PNPLA2 at the neutral lipid-rich monolayer of LDs and PNPLA3 at the phospholipid bilayer of the endoplasmic reticulum (ER) (15). The regulatory impacts of membrane composition on protein dynamics are increasingly recognized as critical determinants of biological function, yet how ABHD5 adapts its conformation to these different lipid environments remains unexplored. Moreover, the reciprocal effects of ABHD5 binding on local membrane structure and lipid reorganization that could influence PNPLA enzyme activity are completely unknown.

Here, we present an integrated multi-scale investigation of ABHD5-lipid interactions that combines computational and experimental approaches to elucidate the structural basis of its regulatory function. Our computational framework employs coarse-grained (CG) (24) and all-atom Gaussian accelerated molecular dynamics (GaMD) (25) simulations to determine ABHD5-membrane orientation and conformational dynamics. These predictions are rigorously validated through site-directed mutagenesis targeting computationally identified key residues (26), live-cell and model membrane fluorescence microscopy to visualize membrane localization dynamics (27), and hydrogen-deuterium exchange mass spectrometry (HDX-MS) to map protein conformational changes upon membrane binding (28). This multidisciplinary approach provides an atomistic-resolution view of ABHD5 structure and dynamics as it interacts with biological membranes. Our findings elucidate the general principles by which ABHD5 regulates lipid metabolism at organelle surfaces, providing a framework for studying membrane-binding proteins with complex, disordered regions. By bridging protein structure prediction and membrane modeling, this study advances our understanding of lipid signaling interfaces and opens new avenues for targeting ABHD5 in metabolic disease.

## Results

ABHD5 can be cytosolic or tightly associated with LD; however, it was unclear to what degree LD binding is an intrinsic property of ABHD5 or might be mediated by interactions with LD proteins, such as perilipins (PLINs) or PNPLAs. Previous work demonstrated that purified ABHD5 binds micelles composed of dodecylphosphocholine, which mimic the membrane phospholipids (21); however, the relative ability to bind defined membranes containing TAG cargo has not been previously investigated. Therefore, we examined the binding of purified ABHD5 to droplet-embedded vesicles (DEVs), a model membrane system that determines protein sorting between phospholipid bilayers and LD monolayers. Notably, we found that ABHD5 strongly sorts to TAG-containing monolayers, suggesting that LD targeting involves interactions with the LD cargo TAGs (**Fig. 1A-C)**.

To understand the molecular basis of ABHD5 membrane targeting, we employed an integrated computational-experimental pipeline combining multiscale MD simulations with orthogonal biochemical validation. We previously analyzed and validated the solution structure of ABHD5 (23) (**Fig. 1D**). Here, we converted the all-atom ABHD5 conformation to CGMD to enable simulating protein diffusion, membrane binding, and large-scale conformational changes over microsecond timescales (**Fig. 1E, F**). These membrane-bound structures were subsequently refined using all-atom Gaussian accelerated molecular dynamics (GaMD) simulations to resolve detailed amino acid-lipid interactions and local membrane remodeling (**Fig. 1G**).

Our multiscale computational approach enables observing atomistic details of lipid-induced protein conformational changes, protein-induced phospholipid remodeling, and the creation of a TAG nanodomain under ABHD5. Critically, we cross-validate these results with fluorescence microscopy, site-directed mutagenesis, and HDX-MS to experimentally map membrane-binding regions and assess conformational dynamics on live cells and model membranes.

### ABHD5 employs a sequential dual-site mechanism for membrane recognition

Our CGMD simulations revealed that ABHD5 interacts with the membrane in two sequential steps: initial anchoring via the N-terminal region (residues 1-43), followed by a second contact through a membrane-binding domain (residues 180-230) within the insertion segment, which includes the lid helix (residues 198-207) that gates the pseudosubstrate pocket (**Fig. 2, S1, S2, Movie 1-2**). This dual-site interaction was consistently observed on both ER and LD models, with the N-terminal region initiating stable lipid anchoring, followed by engagement of the insertion segment. The CGMD simulations also revealed local rearrangements of the membrane-binding region upon lipid association. Because secondary-structure and elastic network restraints were applied, these rearrangements occurred while maintaining the overall structural integrity of ABHD5 and did not represent unrestricted global conformational changes. Notably, the lid region had not previously been implicated in membrane targeting and transitions from proximal to distal of the psuedosubstrate binding pocket upon lipid association by ABHD5.

**Figure 2:**
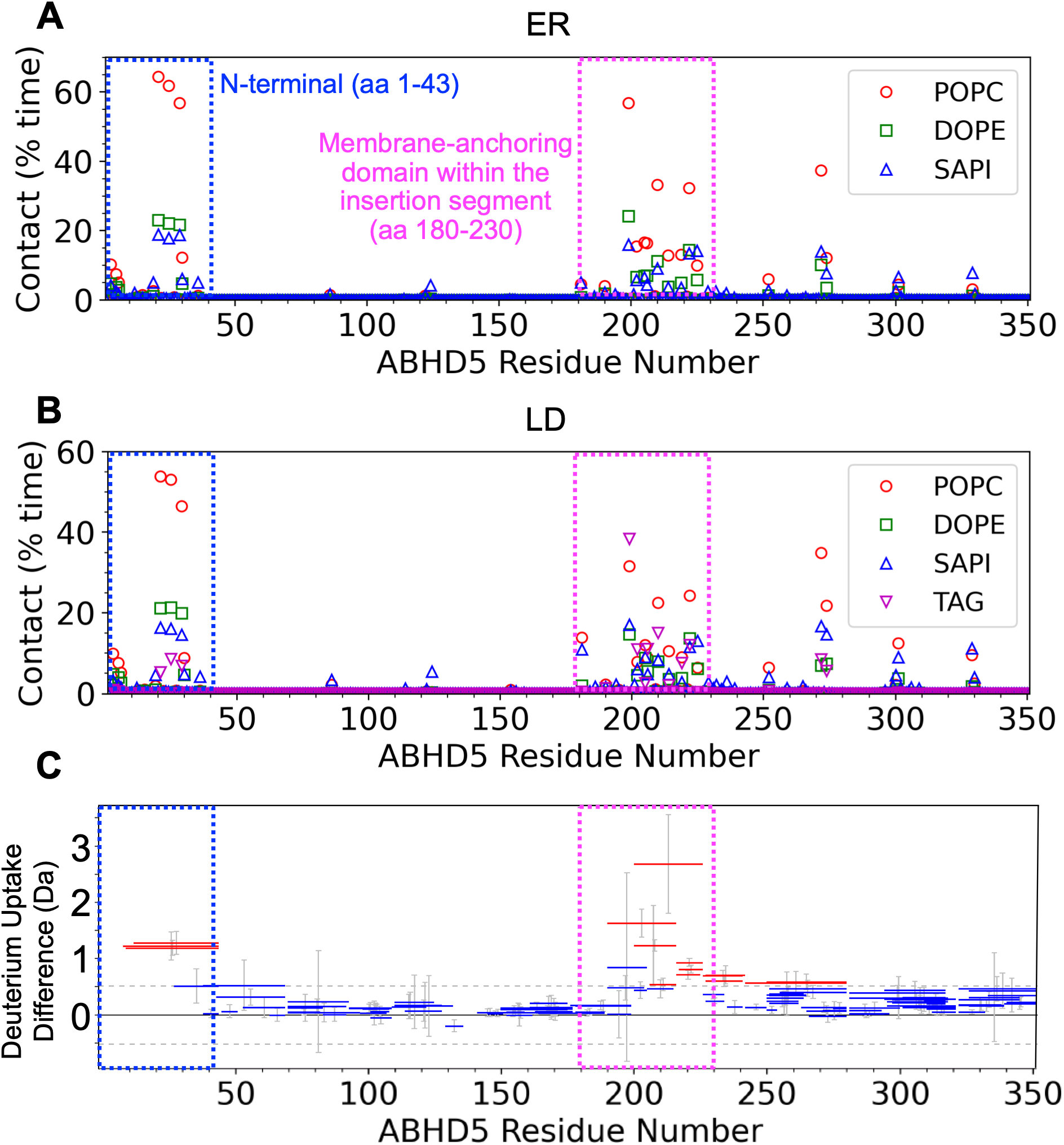
ABHD5 interacts with membranes via two discrete regions. (A-B) Percentage of contact time between each ABHD5 residue and individual lipid species (POPC, DOPE, SAPI, and TAG) from CGMD simulations on ER (A) and LD (B) membranes. Two key membrane-contacting regions are highlighted: the N-terminal region (amino acids 1-43) and a membrane-anchoring domain within the insertion segment (amino acids 180-230). A zoomed view of the interaction pattern in these regions is provided in Figure S1. This data represents the average of contact time from three independent CGMD simulations, with the corresponding error shown in Fig. S2. (C) LD binding induces protection against H/D exchange in the ABHD5 membrane-contacting regions. Horizontal lines indicate peptic peptides from ABHD5 subjected to H/D exchange for 10 s at pH 7.3 and 22°C, in the absence or presence of LDs. The y-axis shows differential deuterium uptake (free ABHD5 minus LD-bound). Fifteen peptides (red lines) spanning residues 8-44 and 190-280 exhibit significant protection upon LD binding. Peptides from other regions (blue lines) show no significant change in deuterium uptake.

To assess lipid specificity, we quantified residue-level contact durations between ABHD5 and individual lipid species. On ER membranes, phosphatidylcholine (POPC) exhibited the most frequent interactions with both N-terminal and lid residues, whereas phosphatidylethanolamine (DOPE) and phosphatidylserine (SAPI) showed rarer contacts (**Fig. 2A, S1A-B**). A similar pattern emerged on LDs, with POPC and TAG interacting robustly with both membrane-binding regions, particularly at the lid domain (**Fig. 2B, S1C-D**).

To independently test our simulation predictions regarding ABHD5-membrane engagement, we employed HDX-MS to test the lipid-dependent changes to the solvent accessibility of ABHD5 peptides. We developed a method to perform HDX on ABHD5 with and without model LD and to remove interfering lipids before MS. Although most peptides of ABHD5 showed no difference in HDX upon membrane binding, we found that the binding of ABHD5 to LD significantly lowered the deuterium uptake of 15 peptides covering two distinct regions: the N-terminal region and the membrane-binding domain of the insertion segment (**Fig. 2C, S3**). The reduced deuterium uptake by ABHD5 when bound to LDs strongly suggests that these regions associate directly with the hydrophobic part of the LDs. The highest difference in deuterium uptake is observed for peptides covering the lid helix within the membrane-binding domain of the insertion segment, with peptide 200-226 having the maximum difference (**Fig. S4**). These experimental results cross-validate our simulation predictions.

### Conserved hydrophobic motifs drive stable lipid interactions and LD selectivity

To characterize the atomic details of ABHD5-membrane interactions, we performed three independent GaMD simulations according to the coordinates from CGMD. The all-atom simulations provided a more detailed description of the interactions underlying the membrane association observed in CGMD. In particular, the N-terminal region consistently penetrated the membrane, with aromatic residues (W21, W25, W29) and hydroxyl-containing residues (T23, T28), forming stable hydrogen bonds and hydrophobic interactions with phospholipid headgroups and tails (**Fig. 3B, E**). The insertion segment includes a lid helix (residues 198-207) composed primarily of hydrophobic residues (V198, W199, I200, A202, V203, G204, A205, A206, L207) whose side chains interact with acyl chains of phospholipid or TAG (**Fig. 3C, F**). The insertion segment also includes additional hydrophobic residues (209–230). The aromatic rings of these residues contribute to membrane association through hydrophobic and aromatic interactions with neighboring residues and lipids.

**Figure 3:**
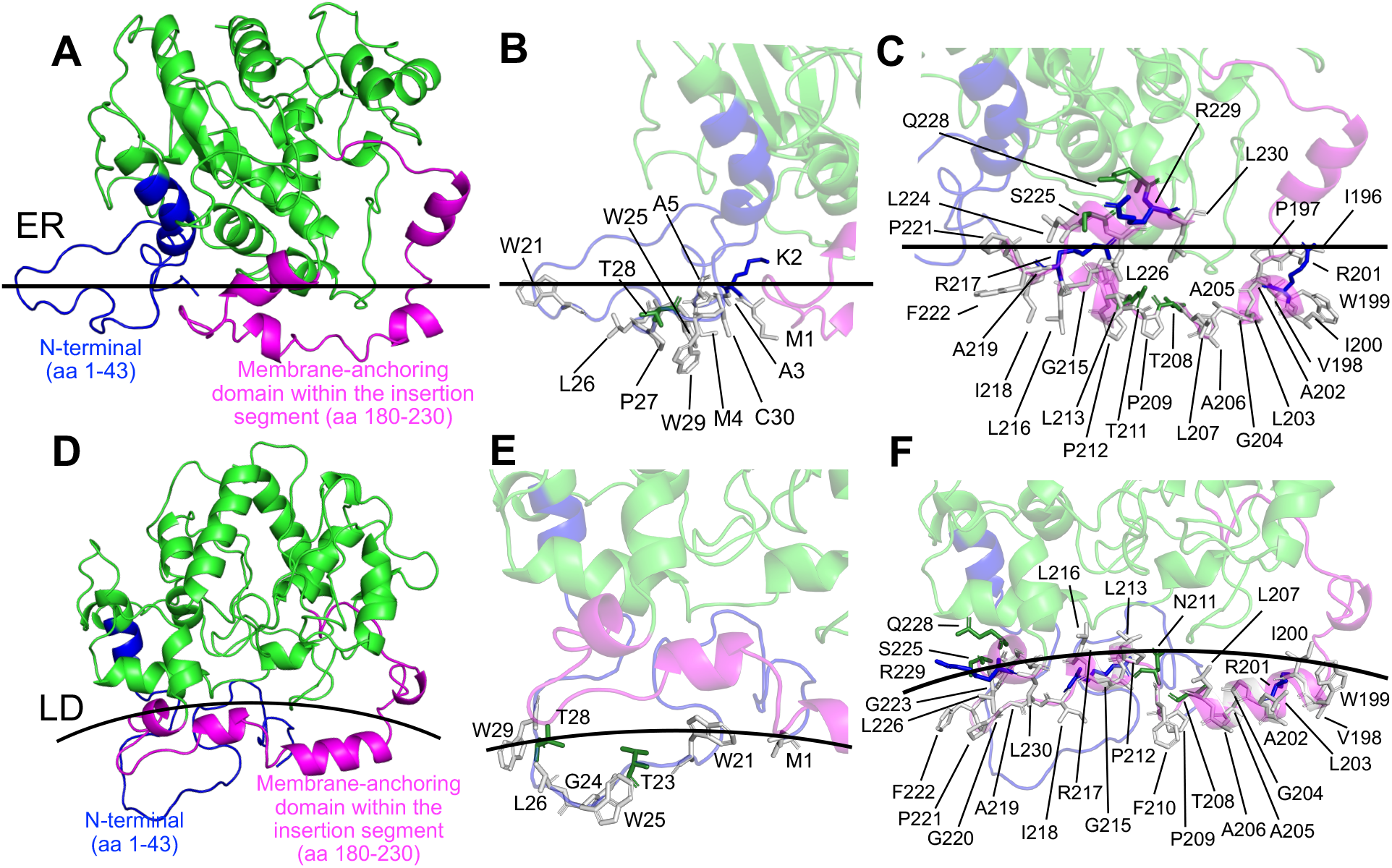
Atomistic simulations reveal insertion of ABHD5 into LD and ER membranes. (A, D) Representative side views from GaMD simulations show ABHD5 inserted into ER (A) and LD (D) membranes. The membrane surface is indicated by a straight (ER) or curved (LD) line. Membrane-interacting regions, N-terminal (residues 1-43, blue) and the domain within the insertion segment (residues 180-230, magenta), are highlighted. (B, E) Close views of the N-terminal interactions with ER (B) and LD (E) membranes. Charged residues are colored blue, polar residues green, and nonpolar residues gray. (C, F) Detailed snapshots of the domain within the insertion segment engaging the ER (C) or LD (F) membrane. Membrane embedding occurs through hydrophobic and electrostatic contacts that differ across organelle-specific lipid environments.

Notably, W199, F210, and F222 form a hydrophobic patch essential for LD association. Both the ER and LD use the same general binding mode for ABHD5 interactions. To experimentally test the significance of these motifs, we mutated these residues to alanine and examined their targeting to LD in transfected Cos7 cells and to model LDs. Wild-type (WT) ABHD5 tagged with cyan fluorescent protein (CFP) was co-transfected with W199A-F210A-F222A ABHD5 tagged with mCherry and imaged using confocal fluorescence microscopy (**Fig. 4A-D**). WT ABHD5 was highly targeted to intracellular LDs of Cos7 cells as evidenced by its strong colocalization with the LD marker BODIPY C16. In contrast, W199A-F210A-F222 ABHD5-mCherry was cytosolic and nucleosolic with negligible association with LDs. Similarly, W21A-W25A-W29A ABHD5-YFP displayed significantly weaker LD targeting than WT ABHD5 (**Fig. S5**). The proper expression and folding of the mutant ABHD5 was confirmed in separate experiments by co-transfecting it with a peptidomimetic PLIN5 (residue 360-417)-YFP and observing the recovery of the mutant ABHD5 LD association through its intact protein-protein binding interface (**Fig. S6**).

**Figure 4:**
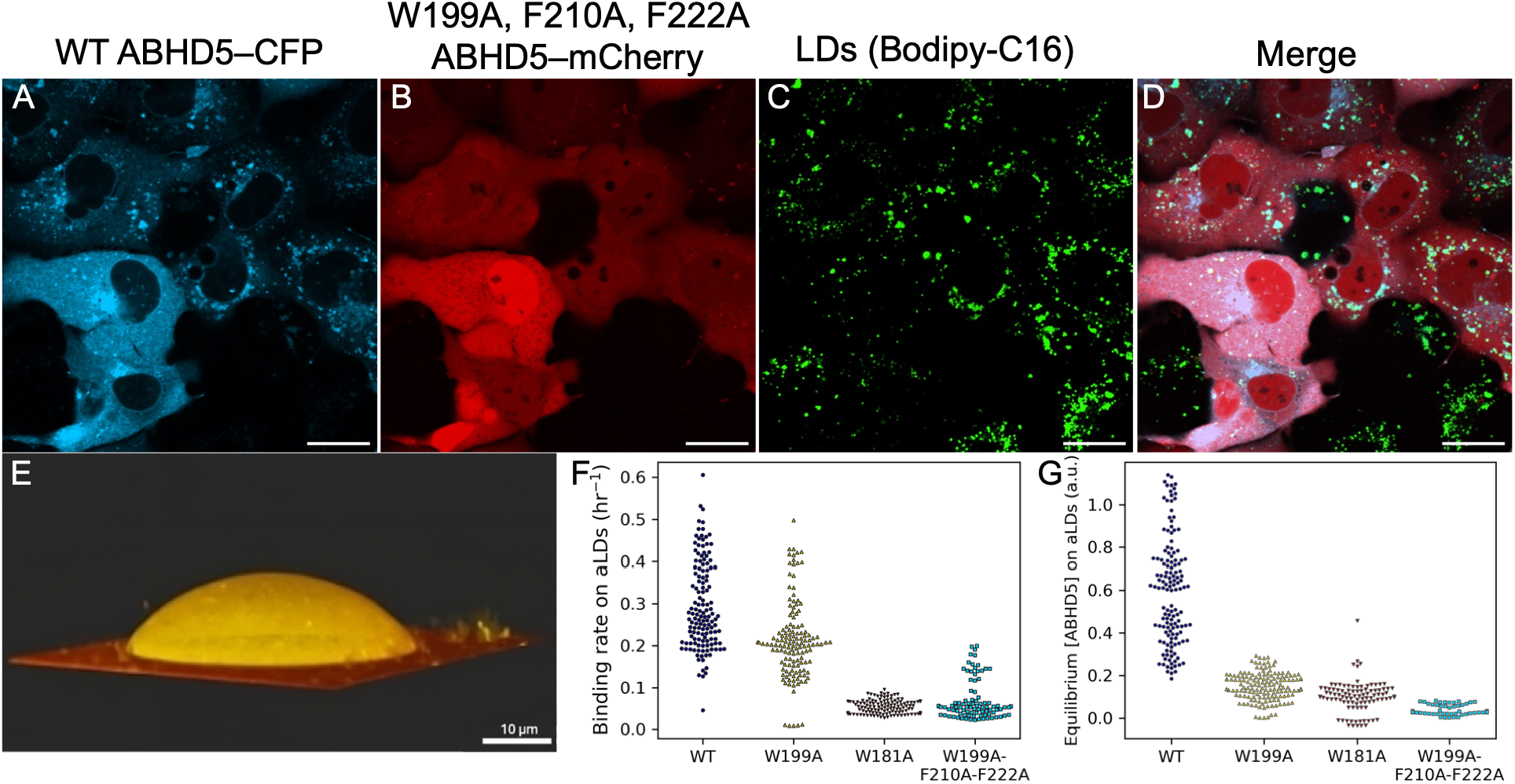
Confocal fluorescence imaging of live cells and model LDs validate the importance of hydrophobic residues for ABHD5 binding to membranes. (A) WT ABHD5 was targeted to LD and ER membranes, whereas (B) the triple-mutant W199A, F210A, F222A ABHD5 was not. (C) LDs were visualized by their incorporation of Bodipy. (D) The color merge image shows the membrane and LD-bound WT ABHD5 colocalized with the LDs while the mutant ABHD5 is cytosolic and within the nucleus. (E) 3D reconstruction image of artificial LD caps shows the concentration of the ABHD5 variants (yellow) on the membrane (red). (F) Mutation of W181A or the triple mutant decreased binding rate of ABHD5 during the first 5 mins to the aLD surface. (G) After 24 hours of equilibration, all three mutants demonstrate significantly lower equilibrium binding to the aLD than the WT ABHD5. (A-D) The scale bars represent 20 µm.

To determine the importance of these key residues on the binding rate and equilibrium binding density of ABHD5, we observed their association with model LDs. Cell lysates expressing WT ABHD5, W199A ABHD5, W181A ABHD5, or W199A-F210A-F222A ABHD5 were exposed to model LDs and imaged with confocal fluorescence microscopy (**Fig. 4E-G**). Mutation of the amino acids predicted to interact with LD lipids strongly disrupted both initial rates and binding at equilibrium. The mutation of W199, which showed the greatest lipid contact in MD, significantly decreased LD binding, which was further significantly decreased when combined with the F210A-F222A mutation. Interestingly, the single W181A mutant was profoundly defective in binding to LDs. MD simulations revealed that W181, while less buried in the hydrophobic core than W199, is positioned at the hinge of the insertion segment and contributes to stabilizing the lid conformation during membrane engagement. These results demonstrate the collective and nuanced contributions of these hydrophobic residues to the mechanism of ABHD5 targeting LDs, including that the central hydrophobic residues contribute more to the equilibrium LD binding density of ABHD5 than the initial binding LD rate.

To evaluate how deeply ABHD5 penetrates the ER and LD membranes, we computed the distance between each residue and the membrane surface. Our analysis revealed that 40 residues of ABHD5 are buried within the membrane for both ER and LD binding, with ABHD5 interacting at similar depths relative to the phosphates of the phospholipid bilayer or monolayer (**Fig. S7**). However, specific contact modes differ between membranes. Residues 285-325 bind deeper within the phospholipids of the LD monolayer than the ER bilayer (22 ± 4 Å vs. 13 ± 7 Å below the phosphates). Notably, R299 was positioned closer to the LD surface than to the ER, with R299 making specific contacts with SAPI. This positioning is functionally significant as R299 is essential for PNPLA2 activation (17, 20), suggesting that membrane-specific positioning facilitates protein-protein interactions critical for lipase activation.

### A lid helix undergoes membrane-induced switching to gate the pseudosubstrate pocket

We hypothesize that the regulation of ABHD5 activity depends on a previously unrecognized conformational switching mechanism involving a cryptic lid helix (residues 198-207) that controls the accessibility of the pseudosubstrate pocket. MD simulations reveal that this hydrophobic helix undergoes dramatic conformational changes upon membrane binding, with the distance between the lid helix and ABHD5 core increasing from 21.8 ± 0.8 Å in solution to 33.3 ± 3.2 Å (ER) and 32.9 ± 1.6 Å (LD) upon membrane engagement (**Fig. 5A-D**). This represents a remarkable ∼53% expansion that transforms the pocket from a closed, water-excluded state to an open, membrane-accessible conformation.

**Figure 5:**
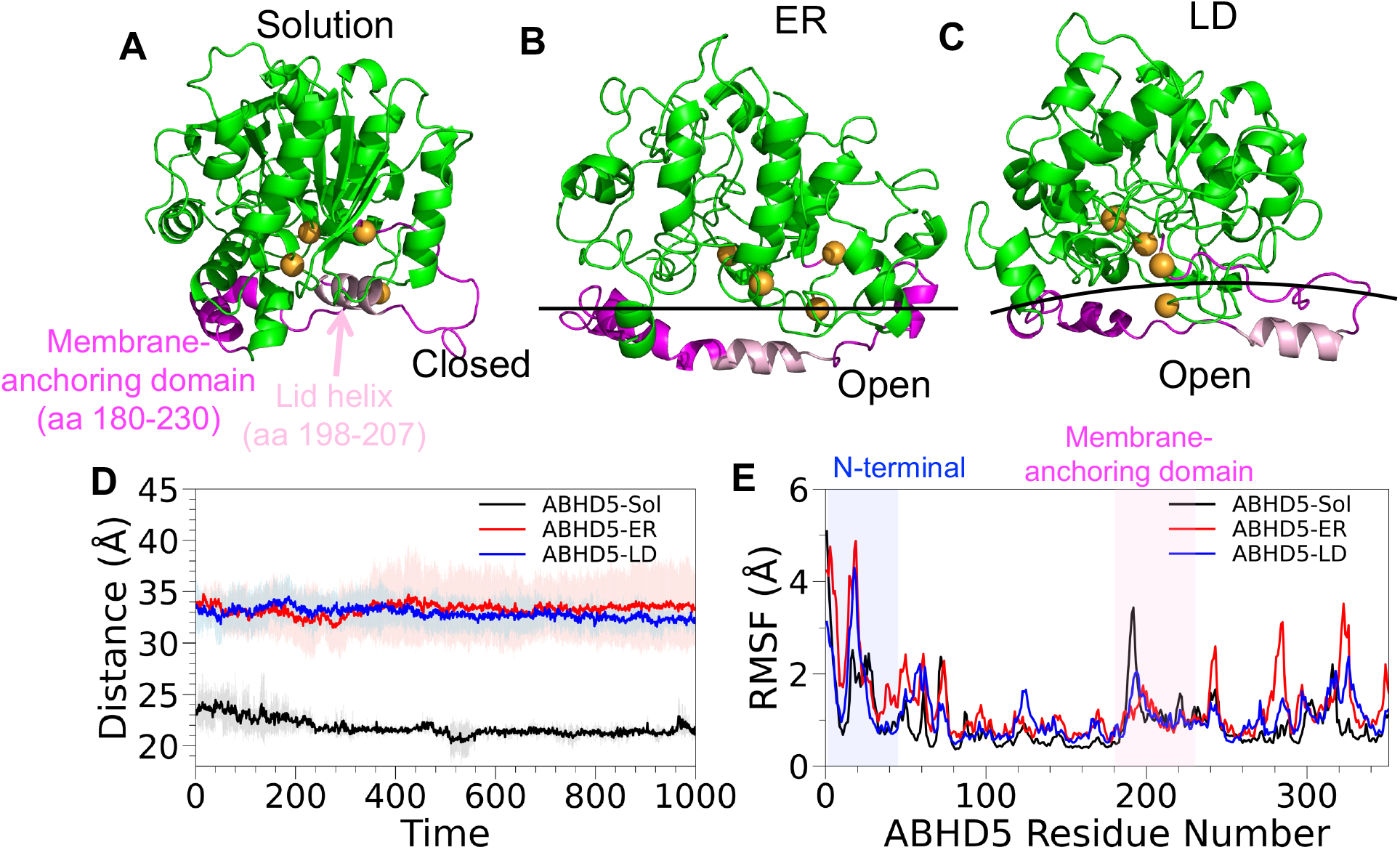
Membrane binding induces conformational changes of the ABHD5 lid helix. (A-C) Representative snapshots from GaMD simulations showing ABHD5 in solution (A), bound to ER membrane (B), and bound to LD membrane (C). The membrane-anchoring domain within the insertion segment (residues 180-230) is shown in magenta; the lid helix (residues 198-207) is highlighted in pink. Key residues within the pseudosubstrate pocket (F86, W181, Y272, Y330) are shown in orange spheres. (D) Distance between the center of mass of the ABHD5 core and the lid helix during GaMD simulations, illustrating lid displacement upon membrane binding. (E) Per-residue Cα RMSF values averaged across three independent simulations for each state (solution, ER-bound, LD-bound). The N-terminal region (residues 1-43) and the membrane-anchoring domain within the insertion segment (residues 180-230) exhibit state-dependent differences in flexibility.

HDX-MS provided orthogonal experimental evidence for the dynamics of the lid. In solution, a peptide (residues 200-226) spanning the lid helix displayed a bimodal isotopic distribution (**Fig. S4**), indicating the coexistence of different conformations. Upon LD binding, the low-exchange population became predominant, demonstrating stabilization of the membrane-bound conformation with reduced solvent accessibility. These experimental observations are consistent with the MD simulations, which reveal lid-helix opening and reduced conformational flexibility upon LD binding. Root-mean-square fluctuation (RMSF) analysis revealed significantly higher conformational flexibility in the lid helix region for the apo state (1.6 ± 0.8 Å) compared to the LD-bound state (1.4 ± 0.4 Å) (**Fig. 5E**). The convergent HDX-MS and MD data establishes a model whereby membrane engagement triggers a conformational transition that stabilizes the open state of the lid helix, facilitating access to the pseudosubstrate-binding pocket.

### ABHD5 remodels local lipid environments to form a TAG-enriched nanodomain

The presence of TAG in LDs alters how ABHD5 interacts with the membrane and reshapes lipid interactions at both its N-terminal and insertion segment interfaces. TAG molecules form contacts with key hydrophobic residues in the insertion segment (W199, A202, A206, F210, and F222) and contacts with N-terminal residues (W21, W25, and W29) (**Fig. 2B, S2D**). The TAG interactions displace POPC from the insertion segment. For instance, POPC interaction with W199 is reduced from 56.7 ± 3.1% in the ER membrane to 31.6 ± 4.2 % in the LD (**Table S1**). However, TAG generally has less effect on DOPE and SAPI interactions.

To determine whether these shifts reflect broader membrane remodeling, we examined the spatial distribution of lipids in the vicinity of ABHD5. On ER membranes, POPC preferentially clusters around the pseudosubstrate pocket, with DOPE displaced and SAPI concentrated under ABHD5 (**Fig. 6A**). In contrast, on the LD, TAG is enriched under the pocket with a sharp perimeter of concentrated SAPI while POPC and DOPE are excluded (**Fig. 6B**). These results suggest that ABHD5 actively reorganized its local lipid environment, displacing phospholipids and concentrating TAG beneath its pseudosubstrate pocket. This localized TAG enrichment may alter the availability and organization of substrate lipids at the ABHD5-LD interface. However, the accessibility of this TAG pool to PNPLA2 and its functional contribution to TAG hydrolysis remain to be established.

**Figure 6:**
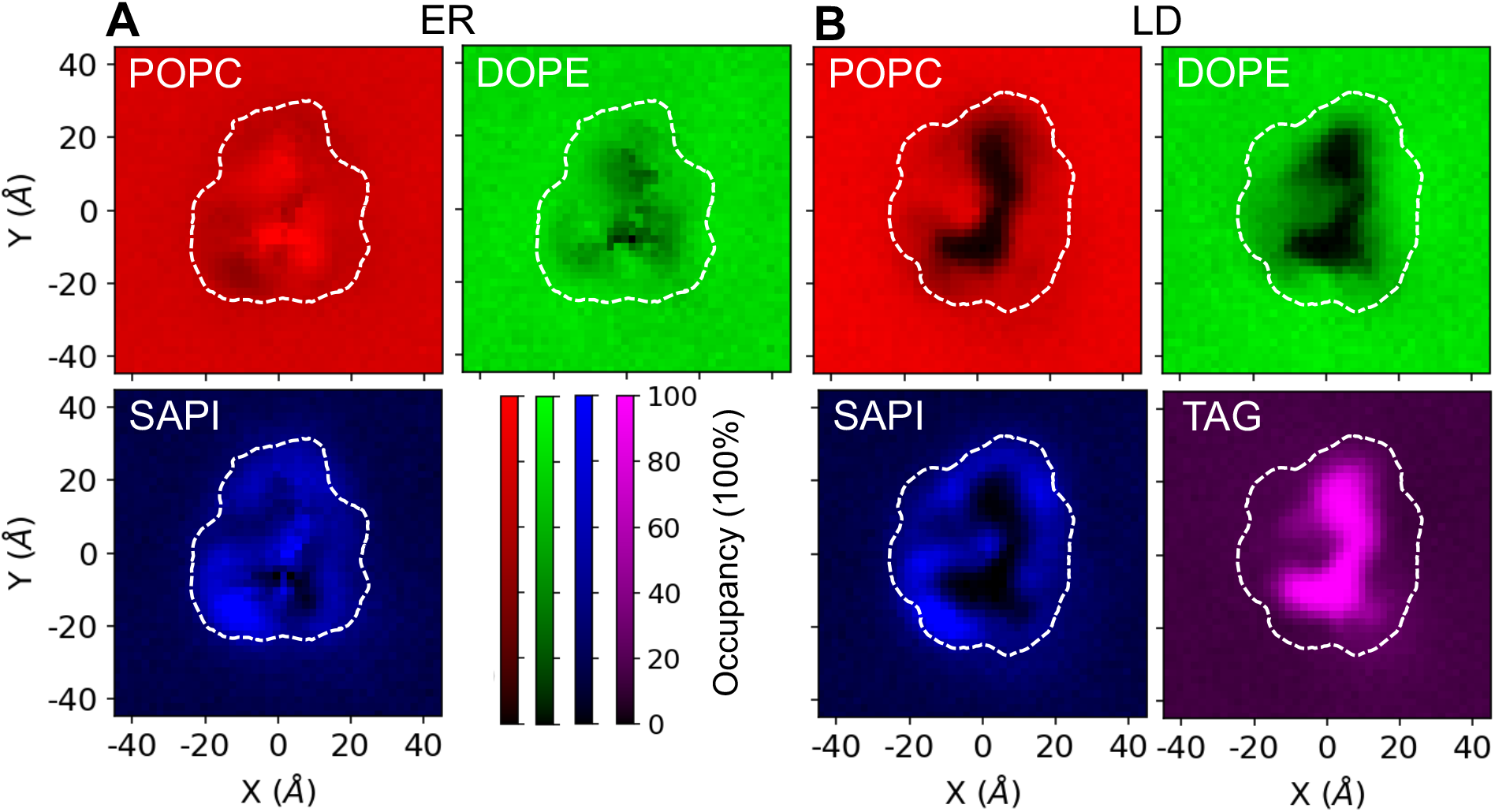
Spatial distribution of lipid species surrounding ABHD5 on ER and LD membranes. Lipid density maps from CGMD simulations showing the average local enrichment of four lipid species, POPC (red), DOPE (green), SAPI (blue), and TAG (purple), around ABHD5 on ER (A) and LD (B) membranes. Darker colors indicate higher local lipid density. The dashed white outlines represent the projected footprint of ABHD5 on the membrane surface.

### ABHD5 induces localized membrane curvature

To investigate whether ABHD5 binding deforms the shape of the phospholipid surface, we calculated membrane surface topology from phospholipid center-of-mass positions and TAG surface topography (**Fig. 7**). Phospholipids were binned across the XY-plane, and mean heights over the final 200 ns of the GaMD simulation defined the average membrane shape for ER and LD systems (**Fig. 7A, B**). On LD membranes, ABHD5 binding induced a pronounced local curvature, elevating phospholipids by 14 ± 5 Å above the surrounding phospholipids. This curvature spanned a lateral distance of 33 ± 3 Å at full width at half maximum (FWHM), though the finite simulation box may underestimate the true extent. In contrast, ER membrane topology displayed no significant change upon ABHD5 binding.

**Figure 7:**
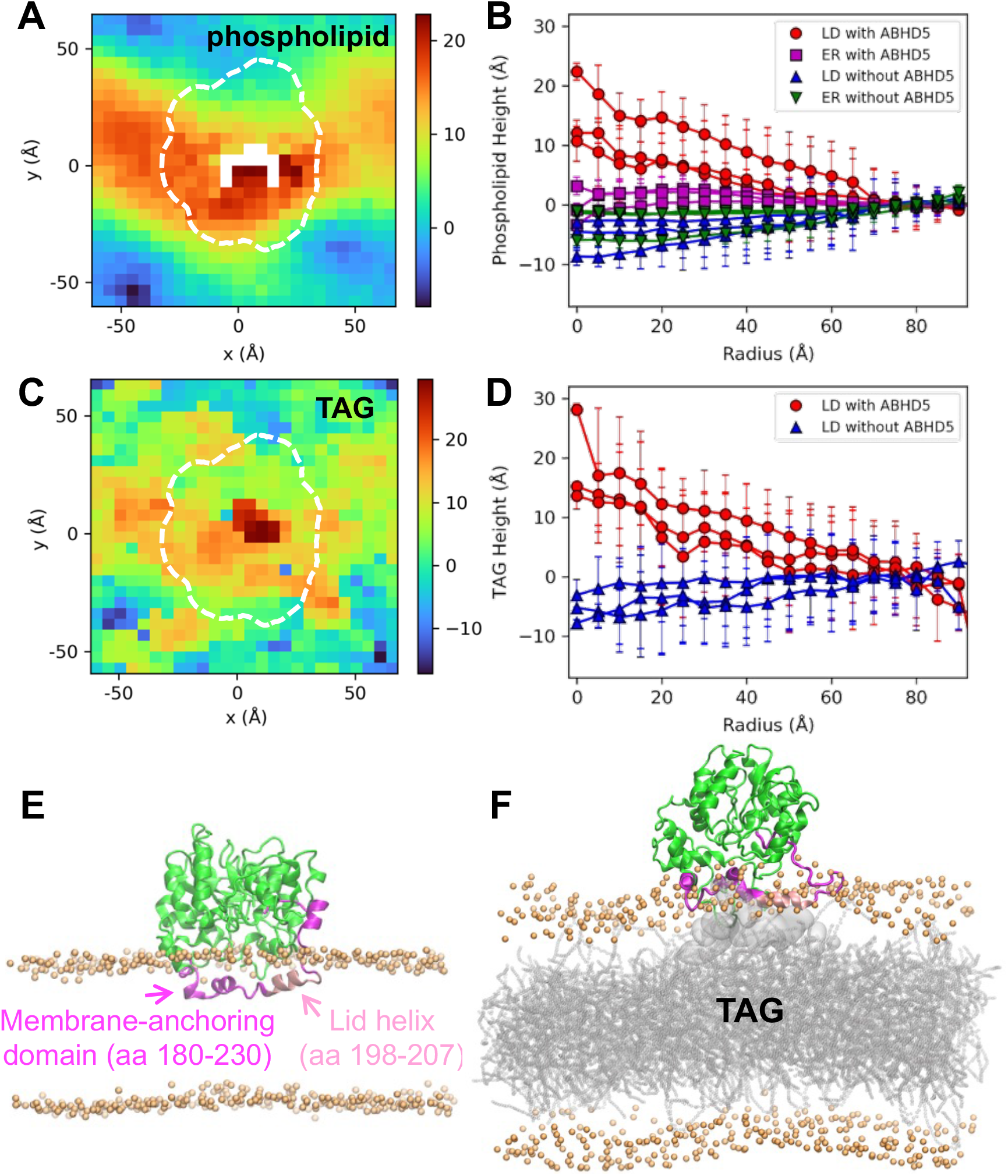
ABHD5 induces local membrane curvature and lipid redistribution on LDs. (A) Time-averaged height map of the LD membrane surface over the final 200 ns of GaMD simulation with ABHD5 bound (dashed outline). A local elevation of phospholipids is observed beneath the protein. White pixels indicate absence of phospholipid coverage. (B) Radially averaged phospholipid height as a function of distance from the ABHD5 center for three independent LD simulations. (C) Time-averaged height map of TAG molecules in the same simulation, showing a pronounced TAG-rich nanodomain beneath ABHD5. (D) Radially averaged TAG height from the same simulation set. Error bars in (B) and (D) represent standard deviations across the 200 ns trajectory window for each replica. (E-F) Side views of ABHD5 bound to ER (E) and LD (F) membranes from representative simulation snapshots. The membrane-anchoring domain (residues 180-230) is highlighted in magenta, and the lid helix (residues 198-207) in pink. POPC, DOPE, and SAPI phosphates are shown as orange spheres; TAG molecules forming the nanodomain are shown as a gray surface/lines.

We next examined whether this remodeling extended to the underlying neutral lipid layer (**Fig. 7D, E**). Analysis of the top TAG surface revealed dramatic remodeling beneath ABHD5. ABHD5 binding generated a distinct, elevated TAG nanodomain (**Fig. 7F, G**) This TAG-rich domain rose 17 ± 5 Å above distal TAG regions and was more prominent than the changes of phospholipids (**Fig. 7C**, **Movie 3-4**). Importantly, the bottom TAG surface and phospholipid monolayer, which lacks bound ABHD5, showed no comparable remodeling (**Fig. S8-S11)**.

## Discussion

ABHD5 plays a central role in lipid metabolism through membrane-restricted activities at the ER and LDs (29–31). While its solution structure has been previously modeled (23, 32), the molecular basis for its membrane association and regulatory function has remained unclear. Here, we establish an integrated computational-experimental framework that reveals, at atomistic resolution, how ABHD5 dynamically engages lipid membranes, undergoes conformational switching, and reshapes its lipid nano-environment. By combining multiscale MD simulations with orthogonal validation via HDX-MS and cell-based assays, we uncover a dual-site membrane recognition mechanism and a cryptic, membrane-responsive regulatory element. This study not only provides new mechanistic insights into ABHD5 function but also introduces a broadly applicable pipeline for dissecting protein-membrane interactions in complex biological contexts.

Our integrated analysis reveals that ABHD5 employs a sophisticated dual-site membrane recognition mechanism (**Fig. 2**), that fundamentally differs from many membrane-binding proteins that rely on a single membrane-anchoring motif, such as an amphipathic helix or hydrophobic loop, to associate with membranes (e.g., BAR, iPLA_2_, or PH domains) (33–35). In ABHD5, the N-terminus initiates membrane association, while the membrane-anchoring domain (residues 198-207) within the insertion segment reinforces and stabilizes the interaction. This two-step process ensures ABHD5 preferentially targets membranes with appropriate lipid compositions, while membranes lacking optimal properties fail to support both binding events. The mutagenesis data show that both binding sites are essential for LD targeting, as disrupting either one greatly reduces it (**Fig. 4** and Ref. 23). This dual-site recognition is critical for ABHD5 cellular function and establishes a new paradigm for organelle-specific protein targeting.

The insertion segment is a defining feature of ABHD5 (**Fig. 1D**). Previous modeling of ABHD5 in solution noted its high flexibility (23), yet its function was not understood. Our findings establish this region as essential for membrane targeting, where it adopts multiple conformations to mediate interaction with both ER and LD membranes. Hydrophobic residues within the insertion segment, including those in the lid helix, penetrate the membrane to engage lipid tails or TAGs, while polar residues interact with lipid headgroups, mimicking the amphipathic helix-based membrane binding used by other peripheral proteins. This binding mode parallels that of proteins such as iPLA₂, which also contains an α/β hydrolase domain and uses an amphipathic helix for bilayer association (36). Similarly, Protein Kinase C (PKC) relies on a diacylglycerol (DAG)- and phosphatidylserine (PS)-responsive amphipathic helix in its C1 domain for membrane anchoring and activation (37). PLIN4 and Arf proteins also employ hydrophobic-rich short helices to interact with monolayers and bilayers (38, 39). The insertion segment of ABHD5, enriched with large nonpolar residues (e.g., valine, tryptophan), enables it to functionally bridge both membrane types. This feature highlights a conserved mechanism among lipid-associated regulatory proteins.

ABHD5 contains a lid helix that undergoes dramatic conformational switching upon membrane binding. The distance between the lid helix and the protein core increases by 1.5-fold (**Fig. 5D**), which transforms ABHD5 from an inactive cytosolic protein into a membrane-localized activator. Interestingly, HDX-MS revealed a bimodal isotopic distribution for the membrane-binding segment in solution (**Fig. S4**), which suggests the coexistence of protected and unprotected conformational states. Further analysis of the exchange profiles supports EX1-like exchange, consistent with dynamic transitions between these states. We propose that the protected state in the absence of LDs represents a closed soluble conformation, whereas in the presence of LDs, it corresponds to an LD-bound conformation. Thus, the bimodality does not necessarily reflect lid-helix formation and unfolding alone, but may reflect broader conformational transitions within the membrane-binding segment. Although LD binding appears to stabilize the membrane-associated protected state, the current data cannot distinguish whether this process involves conformational selection, induced fit, or a combination of both mechanisms. This membrane-induced transition defines a novel regulatory mechanism in which ABHD5 activity is directly governed by lipid binding rather than classical small-molecule allostery. The mechanism is conceptually similar to interfacial activation in lipases, such as Lp-PLA2, where lipid engagement opens access to the catalytic site (40). The large conformational change created by lid opening presents attractive therapeutic opportunities, as small molecule modulators could provide precise control of either open or closed states. Targeting this switch could offer a strategy to modulate ABHD5 activity in metabolic diseases involving dysregulated lipolysis.

The mechanism by which ABHD5, a pseudoenzyme, regulates the activity of PNPLA2, a TAG hydrolase, remains poorly understood. Our findings reveal that ABHD5 binding to LDs induces substantial membrane remodeling, which may influence the local organization and availability of lipid substrates. Upon binding to LDs, ABHD5 induces pronounced local curvature, displaces phospholipids, and concentrates TAG beneath its pseudosubstrate pocket. This spatial reorganization creates a distinct lipid nanodomain at the ABHD5-LD interface. Membrane remodeling is a known strategy for regulating lipid accessibility, particularly in bilayer systems like the ER, where amphipathic helices and lipid modifiers generate packing defects and curvature to influence membrane dynamics (41–44). Unlike bilayers, LDs possess a monolayer rich in neutral lipids that promote curvature and exhibit unique properties, e.g., larger packing defects (39), which influence protein interactions and lipid metabolism by recruiting the amphipathic helices from proteins such as PLINs and CCTα (45–47). This bidirectional interaction, in which LD binding triggers ABHD5 conformational switching and ABHD5 in turn remodels the local lipid landscape, may influence the organization of the LD surface and the interactions of lipid-associated proteins. However, whether ABHD5-mediated remodeling directly facilitates PNPLA2 recruitment or access to TAG substrates remains to be established. In this regard, differences in the interplay between ABHD5 and LD versus ER membranes likely impact its regulation of PNPLA effectors and the distinct pathways under their control.

Our integrated approach reveals how ABHD5 uses dual-site membrane recognition and lid switching to transform from an inactive cytosolic protein into a membrane-localized activator that actively remodels lipid environments. These mechanisms establish new principles for how LD-associated proteins achieve functional specificity through membrane-dependent regulation. Several directions emerge from this work. ABHD5 interacts with PLIN1 and PLIN5, which suppress lipolysis activation, as well PNPLA 1, 3 and 5, whose functions are poorly understood. Future studies should examine how ABHD5 coordinates with PLIN and PNPLA paralogs on distinct membrane surfaces. Elucidating these interactions will reveal how ABHD5 is directed to distinct subcellular locations to control diverse lipid metabolic processes.

We previously reported that ABHD5 is the target of natural and synthetic ligands that bind the pseudosubstrate pocket to regulate the interaction ABHD5 with PLIN and PNPLA paralogs (17, 23, 32), yet the structural basis of those interactions is not fully understood. Modeling ABHD5-ligand complexes in the appropriate membrane environment may support the development of targeted modulators of lipid metabolism. Moreover, the tissue-specific ABHD5 binding partners and regulated pathways suggest that differences in membrane composition across cell types may fine-tune its regulatory roles, which highlights the need to investigate how lipid environments modulate ABHD5-membrane interactions. Overall, our findings establish ABHD5 as a model for how LD-binding proteins coordinate membrane association, conformational transitions, and lipid remodeling to regulate metabolism. These principles provide molecular targets for therapeutic intervention in metabolic diseases where dysregulated lipolysis contributes to pathology.

## Materials and Methods

### Molecular Models

Although the experimental structure of the full ABHD5 protein is unknown, we obtained its structure from a previous study that modeled the mouse ABHD5 protein using homology modeling and deep learning tools, with experimental validation (23). In this study, we examined two membrane systems: 1) A bilayer system that contains POPC, DOPE, and SAPI lipids with a ratio of 88:37:10 to model the ER membrane, and 2) A trilayer composed of a 4 nm thick TAG layer sandwiched between the two phospholipid monolayers modeled the LD (48). A bilayer membrane (to mimic ER) was constructed using the CHARMM-GUI membrane builder (49). A 4-nm (in the *z* dimension) TAG layer was created using PACKMOL (50), resulting in a final density of 0.9040 g/cm^3^. Both the bilayer and TAG membranes had identical x and y dimensions of 14 nm. The membrane (mimicking LD structure) was assembled by inserting the TAG layer into the bilayer with an additional 1-nm spacing along the z dimension between the TAG and each leaflet of the bilayer. The LD membrane system contains 352 POPC, 148 DOPE, 40 SAPI, and 423 TAG molecules. Each system was solvated with TIP3P water molecules (51), with a layer of 15 Å on the top and underneath the bottom of the membrane. A concentration of 0.15 M KCl was added into all systems.

### CGMD Simulations

CG models for protein, ER and LD were constructed using CHARMM-GUI based on the previously described all-atom models (52, 53). The ionizable side chains of the protein were chosen in their default charge states for pH 7. All CGMD simulations were performed using GROMACS 2020.5 (54). The MARTINI 3.0 force field was used for the protein, lipids, ions, and non-polarizable CG water molecules (55). To maintain the protein secondary and tertiary structure, an elastic network model composed of harmonic restraints was applied for MARTINI 3.0 (55). The initial box dimensions and total number of beads in each system are listed in **Table S2**. All systems were minimized in two steps using the steepest descent algorithm: 5000 steps were performed initially, followed by an additional 5000 steps (56). We conducted five equilibration processes with varying lipid restraints and integration time steps. Initially, restraints of 200 kJ/mol/Å^2^ were applied to the lipids with a 2 fs integration time step. Subsequently, the restraints decreased to 100, 50, 20, and 10 kJ/mol/Å^2^with integration time steps of 5, 10, and 20 fs, respectively. During the production simulation, no restraints were applied to the lipids, and the integration timestep remained at 20 fs. The detailed steps are listed in **Table S2**. Temperature was kept constant at 303 K using the velocity rescaling thermostat (57). A Berendsen barostat was employed to implement surface-tension pressure coupling (58), with a coupling constant of 4 ps, a compressibility value of 3×10^−4^ bar^−1^ for both x and y dimension, and a reference pressure of 1 bar. All the simulations were performed with periodic boundary conditions to simulate the bulk behavior. The relative dielectric constant was set to 15 for the system with nonpolarizable water. The long-range electrostatic Coulombic interactions were truncated with a shift cutoff of 1.1 nm. The van der Waals interactions were also smoothly shifted to zero between 0.9 and 1.1 nm. Each simulation was conducted for 20 µs and replicated three times.

### Conventional all-atom MD Simulations

We used Amber20 package for setting up all-atom MD simulations and performing production simulations with an efficient GPU implementation (59). The protein was parameterized using the Amber ff14SB force field (60), while the POPC and DOPE lipids were parametrized using the Lipid14 force filed (61). For the TAG and SAPI lipids, the General Amber Force Field (GAFF) was employed (62). All systems were solvated using the TIP3P water model (51). A salt concentration of 0.15 M KCl was added to the systems using the ion parameters developed by Joung and Cheatham (63). All systems underwent two processes of minimization, first with 2000-5000 steps of the steepest descent method (56), followed by 3000-5000 steps of the conjugate gradient algorithm (64). After minimization, the systems were heated through two processes with fixed lipids. Initially, we increased the temperature from 0 to 100 K over 10 ps, followed by heating from 100 to 303 K over 200 ps. Subsequently, the entire system was equilibrated without restraints through ten repeated simulations, each lasting 1 ns at a temperature of 303 K. Before conducting enhanced sampling, the 50 ns conventional MD production simulations were performed after equilibration. The anisotropic Berendsen weak-coupling barostat was employed to maintain pressure (58), while the temperature was regulated using the Langevin thermostat with a collision frequency of 5 ps^−1^ (65). Bonds containing hydrogen atoms were restrained using the SHAKE algorithm (66). The long-range electrostatics were accounted for using the particle mesh Ewald summation method with a cutoff of 12 Å (67). The time step of the simulation was set to 2 fs.

### GaMD Simulations

GaMD is a powerful enhanced sampling method by adding a harmonic boost potential to smooth the potential energy surface of a biosystem (25). GaMD enhances the efficiency of exploring the energy landscape without constraints (68). Our GaMD simulations comprised a 60-70 ns preparation phase, followed by a production run of 1000 ns. The preparation phase commenced with 8-12 ns (ntcmd) of conventional MD at 303 K, used to gather potential energy statistics necessary for calculating the harmonic force constant and threshold energy parameters essential for the boost potential evaluation. Next, an 4-6 ns GaMD simulation was performed with applied boost potential (ntebprep), but without updating the maximum and minimum potential values. Then, GaMD simulation was carried out with the boost potential updated over 50 ns (nteb). Finally, we performed 1000 ns production GaMD simulation with a fixed boost potential for all systems. Details are addressed in **Table S3**. The threshold energy was set to the lower bound for all simulations (iE=1). The average and standard deviation of the system potential energies were calculated every 500 ps. The boost potential was added to both the dihedral energy and the total potential energy (igamd=3). The upper limit of the standard deviation for both the total potential energy (sigma0P=6) and the dihedral energy (sigma0D=6) was set to 6.0 kcal/mol. All GaMD simulations were conducted in the NPT ensemble. The setting of Langevin dynamics (65), particle mesh (67), hydrogen restraint (66), cutoff, and timestep are consistent with those used in conventional MD. We collected the resulting trajectories every 10 ps for subsequent analysis.

### LD Membrane Simulations

We constructed an all-atom model for the LD membrane system using CHARMM-GUI and PACKMOL as described previously. Subsequently, we employed conventional MD, GaMD, and CGMD simulations to equilibrate the LD system. Details can be found in **Table S3**. Production simulations were performed for 1000 ns. The mass density of each lipid was calculated to verify the system had reached an equilibrated state (**Fig. S12**). Following that, we employed this equilibrated LD system for further investigation involving the ABHD5 protein.

### ABHD5-membrane simulations

We conducted diffusion simulations using CGMD to investigate the binding mechanism of the full ABHD5 protein to both ER and LD membranes (**Fig. 1E**). The pre-equilibrated LD membrane with a CG model was utilized. Initially, the ABHD5 protein was positioned 7.5 nm away from the membrane surface in the solvent. Each simulation lasted for 20 µs and was replicated three times. To maintain the overall structural integrity of ABHD5 during CGMD, tertiary-structure restraints based on an elastic network were applied. These restraints limited large-scale structural rearrangements while allowing local structural adjustments associated with membrane engagement. Thus, the CGMD simulations primarily captured ABHD5 membrane association, protein orientation, membrane insertion, and rearrangement of the membrane-binding regions, rather than unrestricted global conformational changes. The CGMD simulations revealed initial membrane targeting by the N-terminal region, followed by the rearrangement of the insertion segment, including the lid region, and membrane remodeling. Based on the CG model of the ABHD5-membrane bound conformation (**Fig. 1F**), we used the Multiscale Simulation Tool to convert the CG to all-atom models (69). This tool requires all-atom to CG mapping and isomeric information (i.e., cis/trans/dihedral/chiral). The conversion process from CG to all-atom model involves two steps: 1) randomly placing all-atom atoms near the corresponding CG beads according to provided mapping scheme, and 2) conducting energy minimization with a modified all-atom force field using the OpenMM software (70). Then, we applied the MD protocol as described previously to conduct minimization, equilibration, conventional MD, and GaMD simulations (see **Table S3**). The all-atom GaMD simulations were used to investigate protein-lipid interactions, membrane insertion, and membrane-associated conformational transitions at atomic resolution. Production GaMD simulations were executed for 1000 ns and replicated three times. The three independent replicas yielded similar, although not identical, membrane-bound conformations of ABHD5. Quantitative analyses were performed across the three replicas, and results are reported as mean ± standard deviation unless otherwise indicated.

### MD Trajectory Analysis

We used visual molecular dynamics (VMD) (71) and PyMOL (72) software to visualize the systems and create figures and movies. The analysis of GaMD simulation trajectories included root mean squared fluctuation (RMSF), hydrogen bond analysis, lipid mass density profiles, which were calculated using VMD and the CPPTRAJ program (73). We analyzed CG simulation trajectories using *gmx* in GROMACS (54). The lipid distribution around ABHD5 was computed using the Python scripts available in the Shared Data. The percentage of contact time of each ABHD5 residue with lipid components (POPC, DOPE, SAPI, and TAG) were calculated using the MDAnalysis tool (74). The reported values are averages from three independent simulation runs. We measured the distance between each ABHD5 residue and the membrane surface by calculating the distance along the z-axis between the alpha carbon (Ca) of each residue and the center of mass of the phosphorus atoms within 10 Å of each residue’s x- and y-coordinates on the membrane surface. To assess how the lid helix regulates the opening and closing of the pseudosubstrate pocket, we measured the distance between the ABHD5 protein core region and the lid helix. The ABHD5 core consists of six α-helices and eight β-strands, defined as follows: β1 (residues 53–59), β2 (residues 64–71), β3 (residues 80–84), αA (residues 89–103), β4 (residues 106–111), αB (residues 126–143), β5 (residues 149–154), αC (residues 157–171), β6 (residues 172–181), αD (residues 278–288), β7 (residues 292–298), αE (residues 305–314), β8 (residues 319–325), and αF (residues 331–351). The lid helix includes residues 198-207. To evaluate the impact of ABHD5 binding on membrane, we analyzed the membrane surface shape by tracking the center of mass positions of each lipid. The lipids were grouped into bins across the XY-plane, and the average lipid height in each bin was determined over the last 200 ns of the GaMD simulation.

### Analysis of ABHD5 targeting in Cos7 cells and model membranes

WT and mutant ABHD5 were fused to the N-terminus of mCherry as previously described (23). W199A-F210A-F222A mutations were generated by PCR, and all mutations were verified by DNA sequencing. Cos7 cells were plated onto 25 mm glass coverslips and transfected with plasmids encoding WT or mutant ABHD5. In some experiments, cells were co-transfected with PLIN5-EYFP, an LD scaffolding protein that binds ABHD5 (75), or LiveDrop-ECFP, a maker of LDs (76). Following transfection, cells were incubated in 200 μM oleic acid complexed to BSA to facilitate the development of intracellular LD (75). After overnight incubation cells were fixed in 4% paraformaldehyde and imaged using a Dragonfly spinning disc confocal microscope and 63X 1.4 NA oil objective. Human embryonic kidney 293A cells were transfected with plasmids encoding mCherry-tagged WT or mutant ABHD5, and 18-24 hours later, cells were collected for preparation of cell extracts. Cells from two 10 cm plates were resuspended in 700 μL of intracellular buffer, then disrupted by passing through a 26-gauge needle 10 times, followed by brief sonication. Lysates prepared were centrifuged at 14,000 x g for 5 minutes, and the supernatant was collected for fluorescence determination.

Model LD samples were created as described previously (77). Briefly, Triolein (Nu-Chek Prep Inc.) was mixed with 0.195 w/w% DOPC, 0.081 w/w% DOPE, 0.024 w/w% liver-PI, and 0.003 w/w% TF-PC (Avanti Polar Lipids) at a concentration of 0.3 w/w%. 3 μL of the triolein and phospholipid mixture was added to each well of a coverslip-bottom 96-well plate (Cellvis) and dispersed into micron-sized droplets on the coverslip using a nitrogen stream. The coverslips were coated with intracellular buffer and cell lysates containing the fluorescently labeled WT or mutant ABHD5 diluted to a consistent ABHD5 concentration. The intensity of ABHD5 binding to the surface of the model LD over time was observed for 24 hours using a Dragonfly spinning disc confocal microscope with a 63x 1.4 NA oil-immersion objective and quantified using ImageJ.

Droplet-embedded vesicles were prepared by exposing giant unilamellar vesicles (GUVs) to a triolein emulsion, as previously described (78). GUVs were made through electroformation in a growth chamber with 180 μg of the LD-mimicking PC:PE:PI phospholipid mixture with 0.3 mol % Cy5 dried for >30 min under vacuum on ITO plates prior to filling with 0.75 mL of 200 mM sucrose, and subjected to a 10 Hz, 4 Vpp sine wave function generator for 3 h at 50°C. TAG droplets and GUVs were diluted for osmotic balance in a 66% intracellular buffer and 33% Milli-Q water mixture. TAG droplets were created by mixing 5 μL of >99% pure glyceryl trioleate (T-235, NuChek Prep) with 70 μL of diluted IB, vortexing for 60 s, and bath sonicating for 10 s until a cloudy mixture was observed. Droplet-embedded vesicles were created by mixing 10 μL of GUVs with 20 μL of the TAG emulsion and storing them on the rotator for 30 min at RT. 5 μL of the droplet-embedded vesicles were mixed with 40 μL of diluted intracellular buffer and 100 nM of fluorescence-normalized cell lysates. The droplet-embedded vesicles were imaged in a glass-bottom 394-well plate on a Dragonfly confocal microscope with a 63x 1.4 NA oil-immersion objective.

### HDX-MS

LD-bound ABHD5 was prepared by incubating ABHD5 (185 nM) with 0.74% (v/v) LDs in intracellular buffer (10 mM HEPES, 140 mM KCl, 6 mM NaCl, 1 mM MgCl_2_, pH= 7.3) for 30 min at 22°C. H/D exchange of LD-bound ABHD5 and free ABHD5 was initiated by a two-fold dilution into deuterated intracellular buffer (pH = 7.3, uncorrected pH meter reading) yielding a 50% (v/v) D_2_O content in the exchange solution. Isotopic exchange of ABHD5 (93 nM, 50 pmol per sample) occurred for 10, 100, and 1000 s at 22°C (n=3). After the given labeling time the exchange reaction was quenched by acidification by adding concentrated formic acid to a final concentration of 0.5% (v/v) yielding pH 2.5. Subsequently, in solution digestion at quench conditions was carried out with pepsin at an E:S ratio of 1:1 (w:w), while centrifugating the quenched solution at 20200 RCF at 1°C for 3 min to separate the LDs, which have a lower density, from the bulk solution. To avoid injecting LDs into the LC, the lower 90% of the sample was taken up and snap-frozen in liquid N_2_. The samples were stored at −70 °C until LC-MS analyses.

The LC-MS analysis of all labeled samples and controls was done using a cooled nanoACQUITY UPLC HDX manager system (Waters Corporation, MA, USA) coupled to a Synapt G2 mass spectrometer (Waters Corporation, MA, USA) as described previously (79). Briefly, the quenched samples were thawed at room temperature by placing them in a micro centrifuge for 3 min and subsequently injected into a 700 µl sample loop. The peptides were trapped on a precolumn, then they were separated by reverse phase chromatography and eluted into the electrospray ionization (ESI) ion source and were analyzed by the Synapt G2 quadrupole time-of-flight tri-wave ion mobility mass spectrometer (Waters Corporation, MA, USA).

The raw data from the HDX experiment was compared against a peptide list with masses and retention times for ABHD5 peptic peptides identified from a LC-MS/MS experiment using an Orbitrap Eclipse mass spectrometer (Thermo Scientific). The data was analyzed by DynamX 3.0.0. (Waters) to find the deuterium uptake for all peptides. Statistical significance was evaluated by a hybrid statistical analysis approach based on a global ΔHX significance threshold combined with Welch’s t-tests developed by Weis and coworkers (80). In our experiments, ΔHX was ±0.516 on a 0.05 significance level. For additional HDX-MS methods, see **Text S1**.

## Data, Materials, and Software Availability

The simulation input files, resulting trajectories, and post-MD analysis scripts are freely available at https://doi.org/10.5281/zenodo.15122721. The mass spectrometry proteomics data have been deposited to the ProteomeXchange Consortium via the PRIDE partner repository with the dataset identifier PXD086029. All other data are included in the manuscript and/or supporting information.

## Acknowledgments

We thank the Wayne State University High Performance Computing Center for providing resources for CGMD and GaMD simulations. This work was supported by WSU startup funds and NIH grant R35GM160192 (to Y.M.H.), the WSU Postdoctoral Fellows Grant (to C.V.K., J.G.G., and Y.M.H.), NIH grants R01DK076629 (to J.G.G. and C.V.K.) and R01DK105963 (to J.G.G.), and the WSU Barber Integrative Metabolic Research Program.

## Author Contributions

Conceptualization: Y.M.H,, C.V.K, and J.G.G. Project design: A.K. and Y.M.H. Investigation: A.K. performed all simulations. A.K., C.V.K, and Y.M.H. analyzed data. Experimental validation: mutagenesis and imaging experiments performed by M.S., H.Z., L.Z., S.P., C.V.K, and J.G.G.. Hydrogen-deuterium exchange experiments performed by M.C.J. and T.J.D.J.. Manuscript writing: A.K., M.C.J., T.J.D.J, C.V.K, J.G.G., and Y.M.H.

## Movie Legends

**Movie 1: Modeling the diffusion of ABHD5 to the ER using CGMD.** All color representations are consistent with those in Figure 1.

**Movie 2: Modeling the diffusion of ABHD5 to the LD using CGMD.** All color representations are consistent with those in Figure 1.

**Movie 3: Modeling the dynamics of ABHD5 bound to the ER using all-atom GaMD.** All color representations are consistent with those in Figure 7.

**Movie 4: Modeling the dynamics of ABHD5 bound to the LD using all-atom GaMD.** All color representations are consistent with those in Figure 7.

## References

1. H. Sunshine, M. L. Iruela-Arispe, Membrane lipids and cell signaling. Curr. Opin. Lipidol. 28, 408–413 (2017).

2. K. Liu, M. J. Czaja, Regulation of lipid stores and metabolism by lipophagy. Cell Death Differ. 20, 3–11 (2013).

3. F. Khan, et al., Unraveling the intricate relationship between lipid metabolism and oncogenic signaling pathways. Front. Cell Dev. Biol. 12, 1399065 (2024).

4. A. L. Santos, G. Preta, Lipids in the cell: organisation regulates function. Cell. Mol. Life Sci. 75, 1909–1927 (2018).

5. C. Boutari, C. S. Mantzoros, A 2022 update on the epidemiology of obesity and a call to action: as its twin COVID-19 pandemic appears to be receding, the obesity and dysmetabolism pandemic continues to rage on. Metabolism 133, 155217 (2022).

6. M. Di Cesare, et al., The heart of the world. Glob. Heart 19 (2024).

7. C. Y. Park, et al., Dysregulation of lipid droplet protein expression in adipose tissues and association with metabolic risk factors in adult females with obesity and type 2 diabetes. J. Nutr. 153, 691–702 (2023).

8. Y.-L. Wu, et al., Epigenetic regulation in metabolic diseases: mechanisms and advances in clinical study. Signal Transduct. Target. Ther. 8, 98 (2023).

9. Z. Wang, et al., Metabolic disorders and risk of cardiovascular diseases: A two-sample mendelian randomization study. BMC Cardiovasc. Disord. 23, 529 (2023).

10. A. Srebrnik, S. Brenner, B. Ilie, G. Messer, Dorfman-Chanarin syndrome: Morphologic studies and presentation of new cases. Am. J. Dermatopathol. 20, 79–85 (1998).

11. M. Schweiger, A. Lass, R. Zimmermann, T. O. Eichmann, R. Zechner, Neutral lipid storage disease: genetic disorders caused by mutations in adipose triglyceride lipase/ *PNPLA2* or *CGI-58* / *ABHD5*. Am. J. Physiol.-Endocrinol. Metab. 297, E289–E296 (2009).

12. Y. Son, et al., Adipocyte-specific Beclin1 deletion impairs lipolysis and mitochondrial integrity in adipose tissue. Mol. Metab. 39, 101005 (2020).

13. Y. Ohno, A. Nara, S. Nakamichi, A. Kihara, Molecular mechanism of the ichthyosis pathology of Chanarin-Dorfman syndrome: Stimulation of PNPLA1-catalyzed ω-O-acylceramide production by ABHD5. J. Dermatol. Sci. 92, 245–253 (2018).

14. T. Hirabayashi, et al., PNPLA1 has a crucial role in skin barrier function by directing acylceramide biosynthesis. Nat. Commun. 8, 14609 (2017).

15. G. Teskey, et al., Lipid droplet targeting of the lipase coactivator ABHD5 and the fatty liver disease-causing variant PNPLA3 I148M is required to promote liver steatosis. J. Biol. Chem. 301, 108186 (2025).

16. A. Lass, et al., Adipose triglyceride lipase-mediated lipolysis of cellular fat stores is activated by CGI-58 and defective in Chanarin-Dorfman Syndrome. Cell Metab. 3, 309–319 (2006).

17. M. A. Sanders, H. Zhang, L. Mladenovic, Y. Y. Tseng, J. G. Granneman, Molecular basis of ABHD5 lipolysis activation. Sci. Rep. 7, 42589 (2017).

18. A. Lass, R. Zimmermann, M. Oberer, R. Zechner, Lipolysis – A highly regulated multi-enzyme complex mediates the catabolism of cellular fat stores. Prog. Lipid Res. 50, 14–27 (2011).

19. S. Romeo, D. B. Savage, Lipase tug of war: PNPLA3 sequesters ABHD5 from ATGL. Nat. Metab. 1, 505–506 (2019).

20. N. Kulminskaya, et al., Unmasking crucial residues in adipose triglyceride lipase for coactivation with comparative gene identification-58. J. Lipid Res. 65, 100491 (2024).

21. A. Boeszoermenyi, et al., Structure of a CGI-58 motif provides the molecular basis of lipid droplet anchoring. J. Biol. Chem. 290, 26361–26372 (2015).

22. A. Gruber, et al., The N-terminal region of comparative gene identification-58 (CGI-58) is important for lipid droplet binding and activation of adipose triglyceride lipase. J. Biol. Chem. 285, 12289–12298 (2010).

23. R. Shahoei, et al., Molecular modeling of ABHD5 structure and ligand recognition. Front. Mol. Biosci. 9, 935375 (2022).

24. S. J. Marrink, H. J. Risselada, S. Yefimov, D. P. Tieleman, A. H. De Vries, The MARTINI Force Field: coarse grained model for biomolecular simulations. J. Phys. Chem. B 111, 7812–7824 (2007).

25. Y. Miao, V. A. Feher, J. A. McCammon, Gaussian accelerated molecular dynamics: Unconstrained enhanced sampling and free energy calculation. J. Chem. Theory Comput. 11, 3584–3595 (2015).

26. “Site-directed mutagenesis” in Methods in Enzymology, (Elsevier, 2013), pp. 241–248.

27. “Fluorescence live cell imaging” in Methods in Cell Biology, (Elsevier, 2014), pp. 77–94.

28. H. Wei, et al., Hydrogen/deuterium exchange mass spectrometry for probing higher order structure of protein therapeutics: methodology and applications. Drug Discov. Today 19, 95–102 (2014).

29. T. Nohara, Y. Ohno, A. Kihara, Impaired production of skin barrier lipid acylceramides and abnormal localization of PNPLA1 due to ichthyosis-causing mutations in PNPLA1. J. Dermatol. Sci. 107, 89–94 (2022).

30. B. Kien, et al., ABHD5 stimulates PNPLA1-mediated ω-O-acylceramide biosynthesis essential for a functional skin permeability barrier. J. Lipid Res. 59, 2360–2367 (2018).

31. Z. H. Jebessa, et al., The lipid-droplet-associated protein ABHD5 protects the heart through proteolysis of HDAC4. Nat. Metab. 1, 1157–1167 (2019).

32. Y. Y. Tseng, et al., Structural and functional insights into ABHD5, a ligand-regulated lipase co-activator. Sci. Rep. 12, 2565 (2022).

33. J. L. Gallop, et al., Mechanism of endophilin N-BAR domain-mediated membrane curvature. EMBO J. 25, 2898–2910 (2006).

34. S. Qin, A. H. Pande, K. N. Nemec, X. He, S. A. Tatulian, Evidence for the regulatory role of the n-terminal helix of secretory phospholipase A2 from studies on native and chimeric proteins. J. Biol. Chem. 280, 36773–36783 (2005).

35. M. A. Lemmon, K. M. Ferguson, J. Schlessinger, PH Domains: Diverse sequences with a common fold recruit signaling molecules to the cell surface. Cell 85, 621–624 (1996).

36. D. Bucher, Y.-H. Hsu, V. D. Mouchlis, E. A. Dennis, J. A. McCammon, Insertion of the Ca2+-Independent Phospholipase A2 into a phospholipid bilayer via coarse-grained and atomistic molecular dynamics simulations. PLoS Comput. Biol. 9, e1003156 (2013).

37. S. S. Katti, et al., Structural anatomy of protein kinase C C1 domain interactions with diacylglycerol and other agonists. Nat. Commun. 13, 2695 (2022).

38. A. Čopič, et al., A giant amphipathic helix from a perilipin that is adapted for coating lipid droplets. Nat. Commun. 9, 1332 (2018).

39. C. Prévost, et al., Mechanism and determinants of amphipathic helix-containing protein targeting to lipid droplets. Dev. Cell 44, 73–86.e4 (2018).

40. V. D. Mouchlis, et al., Lipoprotein-associated phospholipase A_2_ : A paradigm for allosteric regulation by membranes. Proc. Natl. Acad. Sci. 119, e2102953118 (2022).

41. A. Gubas, I. Dikic, ER remodeling via ER-phagy. Mol. Cell 82, 1492–1500 (2022).

42. A.-C. Jacomin, I. Dikic, Membrane remodeling via ubiquitin-mediated pathways. Cell Chem. Biol. 31, 1627–1635 (2024).

43. W. Jang, V. Haucke, ER remodeling via lipid metabolism. Trends Cell Biol. 34, 942–954 (2024).

44. J. Varkey, et al., Membrane curvature induction and tubulation are common features of synucleins and apolipoproteins. J. Biol. Chem. 285, 32486–32493 (2010).

45. N. Krahmer, et al., Phosphatidylcholine synthesis for lipid droplet expansion is mediated by localized activation of CTP:Phosphocholine cytidylyltransferase. Cell Metab. 14, 504–515 (2011).

46. N. Kory, R. V. Farese, T. C. Walther, Targeting Fat: Mechanisms of Protein Localization to Lipid Droplets. Trends Cell Biol. 26, 535–546 (2016).

47. M. Majchrzak, et al., Perilipin membrane integration determines lipid droplet heterogeneity in differentiating adipocytes. Cell Rep. 43, 114093 (2024).

48. S. Kim, J. M. J. Swanson, The surface and hydration properties of lipid droplets. Biophys. J. 119, 1958–1969 (2020).

49. E. L. Wu, et al., CHARMM-GUI membrane builder toward realistic biological membrane simulations. J. Comput. Chem. 35, 1997–2004 (2014).

50. L. Martínez, R. Andrade, E. G. Birgin, J. M. Martínez, PACKMOL: A package for building initial configurations for molecular dynamics simulations. J. Comput. Chem. 30, 2157–2164 (2009).

51. W. L. Jorgensen, J. Chandrasekhar, J. D. Madura, R. W. Impey, M. L. Klein, Comparison of simple potential functions for simulating liquid water. J. Chem. Phys. 79, 926–935 (1983).

52. S. Jo, T. Kim, V. G. Iyer, W. Im, CHARMM-GUI: A web-based graphical user interface for CHARMM. J. Comput. Chem. 29, 1859–1865 (2008).

53. J. Lee, et al., CHARMM-GUI input generator for NAMD, GROMACS, AMBER, OpenMM, and CHARMM/OpenMM simulations using the CHARMM36 additive force field. J. Chem. Theory Comput. 12, 405–413 (2016).

54. D. Van Der Spoel, et al., GROMACS: Fast, flexible, and free. J. Comput. Chem. 26, 1701–1718 (2005).

55. P. C. T. Souza, et al., Martini 3: a general purpose force field for coarse-grained molecular dynamics. Nat. Methods 18, 382–388 (2021).

56. J. C. Meza, Steepest descent. WIREs Comput. Stat. 2, 719–722 (2010).

57. G. Bussi, D. Donadio, M. Parrinello, Canonical sampling through velocity rescaling. J. Chem. Phys. 126, 014101 (2007).

58. H. J. C. Berendsen, J. P. M. Postma, W. F. Van Gunsteren, A. DiNola, J. R. Haak, Molecular dynamics with coupling to an external bath. J. Chem. Phys. 81, 3684–3690 (1984).

59. R. Salomon-Ferrer, A. W. Götz, D. Poole, S. Le Grand, R. C. Walker, Routine microsecond molecular dynamics simulations with AMBER on GPUs. 2. Explicit solvent particle mesh ewald. J. Chem. Theory Comput. 9, 3878–3888 (2013).

60. J. A. Maier, et al., ff14SB: Improving the accuracy of protein side chain and backbone parameters from ff99SB. J. Chem. Theory Comput. 11, 3696–3713 (2015).

61. C. J. Dickson, et al., Lipid14: The amber lipid force field. J. Chem. Theory Comput. 10, 865–879 (2014).

62. J. Wang, R. M. Wolf, J. W. Caldwell, P. A. Kollman, D. A. Case, Development and testing of a general amber force field. J. Comput. Chem. 25, 1157–1174 (2004).

63. I. S. Joung, T. E. Cheatham, Determination of alkali and halide monovalent ion parameters for use in explicitly solvated biomolecular simulations. J. Phys. Chem. B 112, 9020–9041 (2008).

64. S. J. Watowich, E. S. Meyer, R. Hagstrom, R. Josephs, A stable, rapidly converging conjugate gradient method for energy minimization. J. Comput. Chem. 9, 650–661 (1988).

65. J. Liu, D. Li, X. Liu, A simple and accurate algorithm for path integral molecular dynamics with the Langevin thermostat. J. Chem. Phys. 145, 024103 (2016).

66. J.-P. Ryckaert, G. Ciccotti, H. J. C. Berendsen, Numerical integration of the cartesian equations of motion of a system with constraints: molecular dynamics of n-alkanes. J. Comput. Phys. 23, 327–341 (1977).

67. U. Essmann, et al., A smooth particle mesh Ewald method. J. Chem. Phys. 103, 8577–8593 (1995).

68. J. Wang, et al., Gaussian accelerated molecular dynamics: Principles and applications. WIREs Comput. Mol. Sci. 11, e1521 (2021).

69. S. Kim, Backmapping with mapping and isomeric information. J. Phys. Chem. B 127, 10488–10497 (2023).

70. P. Eastman, et al., OpenMM 7: Rapid development of high performance algorithms for molecular dynamics. PLOS Comput. Biol. 13, e1005659 (2017).

71. W. Humphrey, A. Dalke, K. Schulten, VMD: Visual molecular dynamics. J. Mol. Graph. 14, 33–38 (1996).

72. The PyMOL Molecular Graphics System, Version 3.0 Schrödinger, LLC.

73. D. R. Roe, T. E. Cheatham, PTRAJ and CPPTRAJ: Software for processing and analysis of molecular dynamics trajectory data. J. Chem. Theory Comput. 9, 3084–3095 (2013).

74. N. Michaud-Agrawal, E. J. Denning, T. B. Woolf, O. Beckstein, MDAnalysis: A toolkit for the analysis of molecular dynamics simulations. J. Comput. Chem. 32, 2319–2327 (2011).

75. J. G. Granneman, H.-P. H. Moore, E. P. Mottillo, Z. Zhu, Functional interactions between Mldp (LSDP5) and ABHD5 in the control of intracellular lipid accumulation. J. Biol. Chem. 284, 3049–3057 (2009).

76. M.-J. Olarte, et al., Determinants of endoplasmic reticulum-to-lipid droplet protein targeting. Dev. Cell 54, 471–487.e7 (2020).

77. S. A. Gandhi, M. A. Sanders, J. G. Granneman, C. V. Kelly, Four-color fluorescence cross-correlation spectroscopy with one laser and one camera. Biomed. Opt. Express 14, 3812 (2023).

78. S. A. Gandhi, et al., Methods for making and observing model lipid droplets. Cell Rep. Methods 4, 100774 (2024).

79. A. Kumari, et al., ANGPTL3/8 is an atypical unfoldase that regulates intravascular lipolysis by catalyzing unfolding of lipoprotein lipase. Proc. Natl. Acad. Sci. U. S. A. 122, e2420721122 (2025).

80. T. S. Hageman, D. D. Weis, Reliable identification of significant differences in differential hydrogen exchange-mass spectrometry measurements using a hybrid significance testing approach. Anal. Chem. 91, 8008–8016 (2019).

