## Supplementary Information for "Unraveling the Molecular Mechanisms of ABHD5 Membrane Targeting"

**Short Title: ABHD5 Membrane binding**

*Amit Kumar[a], Matthew Sanders[b], Huamei Zhang[b], Li Zhou[b], Shahnaz Parveen[a], Miriam C. Jensen[d], Thomas J.D. Jørgensen[d*], Christopher V. Kelly[a], James G. Granneman[b,c], and Yu-ming M. Huang[a*]*

[a]Department of Physics and Astronomy, Wayne State University, Detroit, MI 48201, USA

[b]Center for Molecular Medicine and Genetics, School of Medicine, Wayne State University, Detroit, MI 48201, United States

[c]Center for Integrative Metabolic and Endocrine Research, School of Medicine, Wayne State University, Detroit, MI 48201, United States

[d]Department of Biochemistry and Molecular Biology, Southern Denmark University, Odense Denmark.

*Corresponding author:

Yu-ming M. Huang

ORCID: 0000-0003-3257-6170

Thomas J.D. Jørgensen

ORCID: 0000-0002-7149-316X

**Text S1: HDX-MS Methods**

The LC-MS analysis of all labeled samples and controls was done using a cooled nanoACQUITY UPLC HDX manager system (Waters Corporation, MA, USA) coupled to a Synapt G2 mass spectrometer (Waters Corporation, MA, USA). The thawed samples were injected into a 700 µl sample loop and then the peptides were trapped and desalted on a ACQUITY UPLC BEH C18 VanGuard precolumn (130 Å, 1.7 µm, 2.1 × 5 mm, Waters) with a flow of 300 µL/min for 5 min.

The peptides were separated by reverse phase chromatography using a ACQUITY UPLC peptide BEH C18 column (1.7 µm, 1.0 × 50 mm, Waters) with a gradient of solvent A (0.23% formic acid) and solvent B (acetonitrile with 0.23% formic acid) at a flow of 40 µl/min. The gradient was as follows: 5-10% solvent B over 0.1 min; 10-50% B over 11.9 min; 50-90% B over 0.1 min; 90% B for 0.1 min; 5% B for 0.1 min. The peptides eluted into the electrospray ionization (ESI) ion source and were analyzed by the Synapt G2 quadrupole time-of-flight tri-wave ion mobility mass spectrometer (Waters Corporation, MA, USA). The peptides made by pepsin digestion of ABHD5 were identified by an LC-MS/MS experiment using an Orbitrap Eclipse mass spectrometer (Thermo Scientific) with data dependent acquisition. Three samples of approx. 150 pmol ABHD5 were loaded into the nanoACQUITY UPLC system with online proteolysis on a column packed with immobilized pepsin and same chromatographic method as in the HDX-MS analysis. The precursor peptide ions in a m/z range of 300-1500 were selected and fragmented to obtain MS/MS spectra.

The raw mass spectrum data from peptide identification of undeuterated samples were searched using MASCOT (matrix science) against a custom-made database containing the sequence of 6xHis-tagged mABHD5 as well as possible contaminating proteins to produce a peptide list with masses and retention times.

Maximally labeled controls ("100% $D_2O$") were made by incubating ABHD5 for 13 h at 37° C in deuterated intracellular buffer, 50% (v/v) $D_2O$ content. The "100% $D_2O$" controls were quenched, centrifugated and digested as described above. "0% $D_2O$" controls were made by directly adding ABHD5 from $H_2O$-intracellular buffer to quenched deuterated intracellular buffer, i.e., ~50% (v/v) $D_2O$ content, 0.5% (v/v) formic acid. "0% $D_2O$" controls were centrifugated and digested as described above. The "100% $D_2O$" and "0% $D_2O$" controls were used to adjust for the artifactual deuterium gain or loss that inadvertently occurs after quench (1). The adjusted values were used for generating the normalized deuterium content displayed in the heatmap. "$H_2O$ controls" of ABHD were made in quenched $H_2O$-intracellular buffer, centrifugated and digested as described above. The concentration of ABHD5 in all controls was 100 nM with 50 pmol/sample.

| Area | Residue | ER (% of contact time) | LD (% of contact time) |
|---|---|---|---|
| N-terminal | W21 | 64.4 ± 0.3 | 53.8 ± 4.4 |
| | W25 | 61.7 ± 1.6 | 53.1 ± 3.6 |
| | W29 | 56.7 ± 4.9 | 46.5 ± 3.5 |
| Insertion Segment | W188 | 4.5 ± 0.0 | 13.8 ± 0.0 |
| | W199 | 56.7 ± 3.1 | 31.6 ± 4.2 |
| | A202 | 15.5 ± 2.2 | 7.7 ± 2.1 |
| | A205 | 16.6 ± 2.7 | 12.0 ± 1.4 |
| | A206 | 16.4 ± 1.9 | 8.7 ± 1.4 |
| | F210 | 33.2 ± 2.3 | 22.5 ± 1.4 |
| | F222 | 32.3 ± 0.6 | 24.3 ± 3.6 |
| | Y274 | 12.1 ± 6.9 | 21.8 ± 2.3 |

**Table S1:** The percentage of contact time from CGMD simulations between ABHD5 residues and POPC in ER and LD membranes. The blue indicates residues where POPC interactions increase in the presence of TAG, while all other residues show a decrease in POPC contact when TAG is present.

| Simulation Protocol | Stage | Simulation steps/time |
|---|---|---|
| System information | $N_{beads}$ | LD: 38715<br>ABHD5 with ER: 34549<br>ABHD5 with LD: 40856 |
| | Box dimension | LD: 137Å x140Å x 260Å<br>ABHD5 with ER: 147Å x 147Å x 224Å<br>ABHD5 with LD: 147Å x 147Å x 260Å |
| System minimization | Minimization | 5000 steps |
| | Minimization | 5000 steps |
| System equilibration | Equ (FC=200) | 10 ns |
| | Equ (FC=100) | 5 ns |
| | Equ (FC=50) | 2 ns |
| | Equ (FC=20) | 1 ns |
| | Equ (FC=10) | 1 ns |
| Production simulation | CGMD | 20 μs |

**Table S2:** Summary of the CGMD simulation protocols and setup. The table outlines the detailed steps involved in CG system preparation and CGMD simulation for LD, ABHD5 with ER, and ABHD5 with LD. $N_{beads}$ indicates the total number of beads in each system. The minimization processes were carried using steepest descent algorithm. Then systems were equilibrated with different force constraints (FC) of 200, 100, 50, 20, and 10 kJ/mol/$Å^2$ at 303 K. The CGMD production simulations were then run for 20 μs.

| Simulation Protocol | Stage | ER | LD | ABHD5 with ER/LD |
|---|---|---|---|---|
| System information | $N_{atoms}$ | 127574 | 343838 | ER: 285416<br>LD: 324145 |
| | Box dimension | 139Å x 136Å x 83Å | 139Å x 136Å x 124Å | ER: 151Å x 151Å x 160Å<br>LD: 151Å x 151Å x 180Å |
| System minimization | $M_{SD}$ | 5000 steps | 5000 steps | 2000 steps |
| | $M_{CG}$ | 5000 steps | 5000 steps | 3000 steps |
| System equilibration | $Equ_{0\text{-}100K}$ | 10 ps | 10 ps | 10 ps |
| | $Equ_{100\text{-}303K}$ | 200 ps | 200 ps | 200 ps |
| | $Equ_{Hold}$ | 1 ns x 10 | 1 ns x 10 | 1 ns x 10 |
| | cMD simulation | 50 ns | 50 ns | 50 ns |
| GaMD preparation | ntcmd | 4 ns | 12 ns | 12 ns |
| | ntebprep | 2 ns | 6 ns | 6 ns |
| | nteb | 50 ns | 50 ns | 50 ns |
| Production simulation | GaMD simulation | 1000 ns | 1000 ns | 1000 ns |

**Table S3:** Summary of AAMD simulation protocols and system setup. This table details the steps involved in preparing the GaMD systems and running production simulations for LD, ER, and ABHD5 in complex with ER/LD. $N_{atoms}$ indicates the total number of atoms in each system. The minimization processes were carried out using steepest descent (SD) algorithm and conjugate gradient (CG) method. GaMD preparation for all systems included three steps: 1) collecting potential energy statistics (ntcmd), 2) carrying out GaMD simulations with a fixed boost potential but without updating it (ntebprep), and 3) performing GaMD simulations with the boost potential actively updated (nteb). The subsequent GaMD production simulations were carried out using the finalized, fixed boost potential.

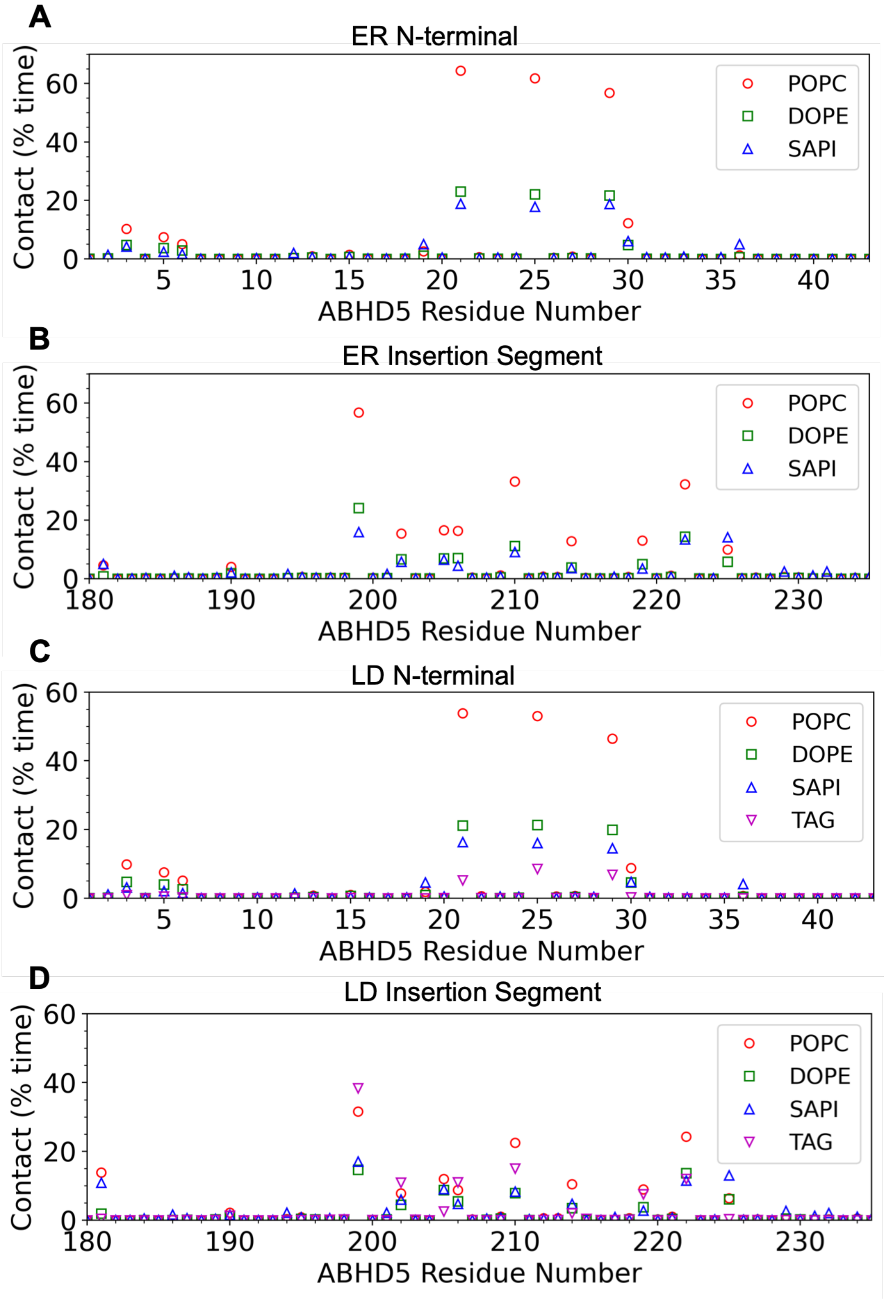


**Figure S1:** The percentage of contact time in CGMD simulations between each ABHD5 residue and various lipid components (POPC, DOPE, SAPI, and TAG) for two key regions — the N-terminal 43 amino acids and the insertion segment (amino acids 180 to 230) — in ER (A, B) and LD (C, D) membranes is shown. These plots provide a zoomed-in view of Figure 2, with the same color scheme used in Figure 2.

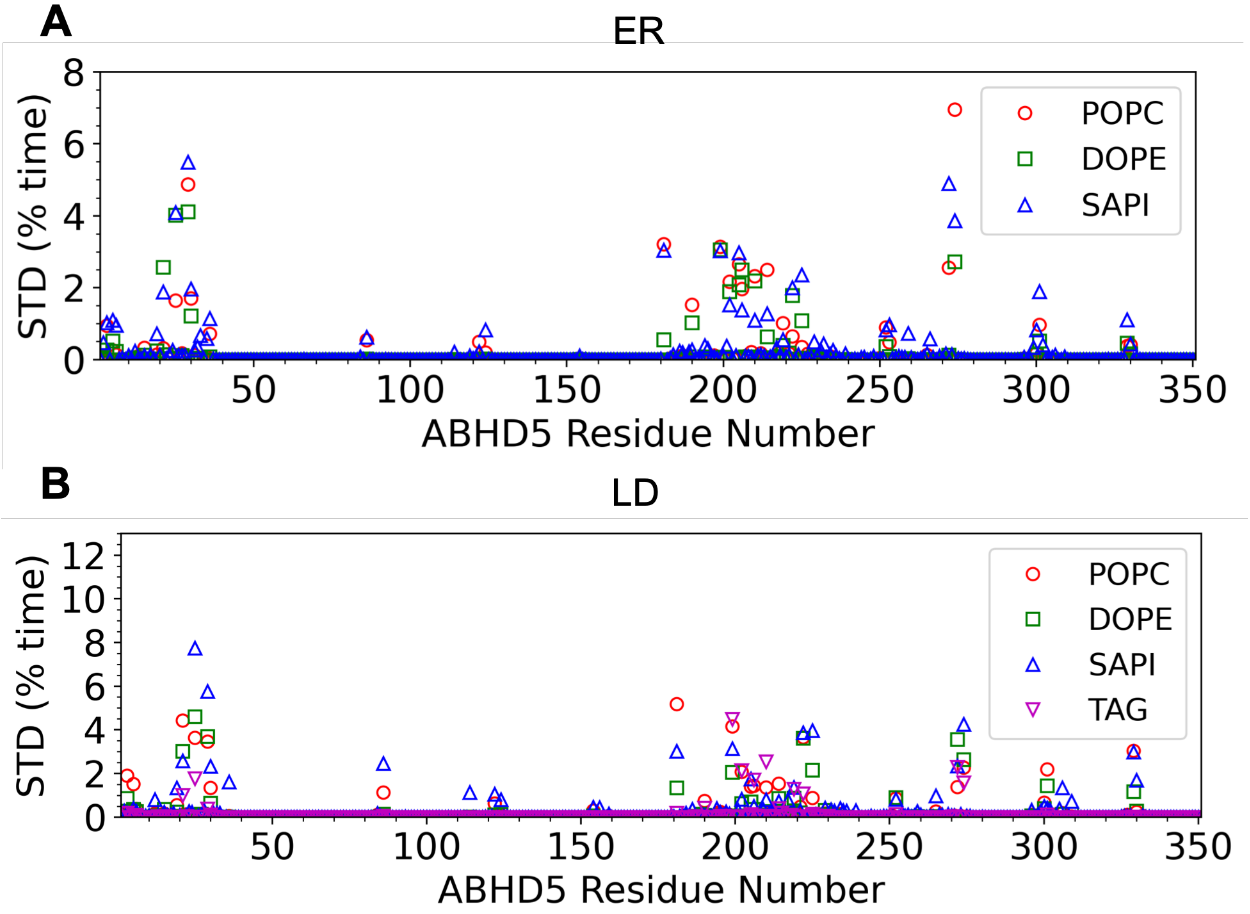


**Figure S2:** The standard deviation of the percentage of contact time from three independent CGMD simulations is shown. The average values are shown in Figure 2, and these plots represent the corresponding standard deviations. The color scheme is consistent with Figure 2 for clarity.

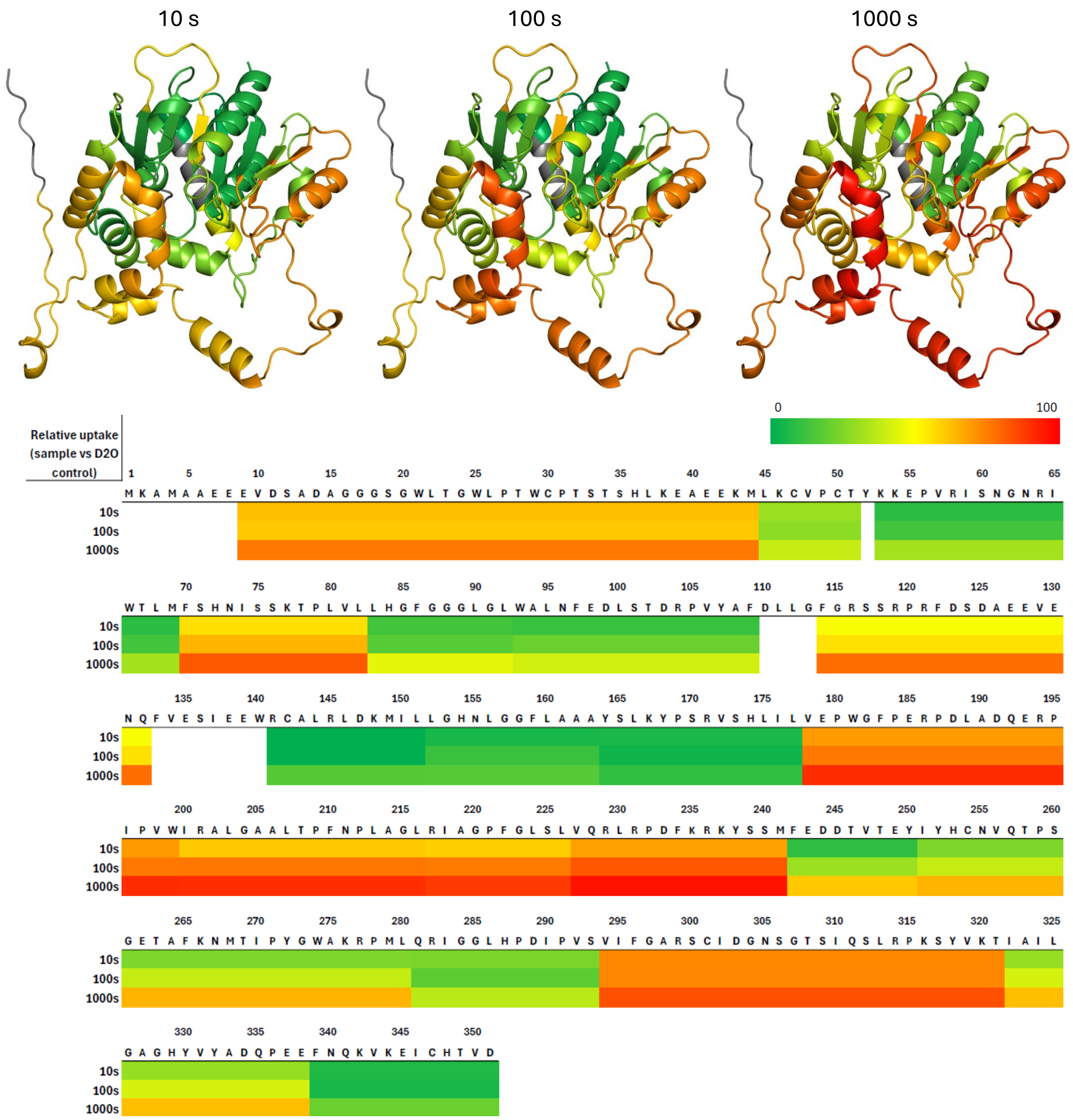


**Figure S3**. Heatmap of the deuterium uptake for selected peptic peptides derived from ABHD5 deuterated without LDs for 10 s, 100 s and 1000 s at pH 7.3, 22°C. The color scale representing deuterium uptake is relative to the 0% and 100% $D_2O$ controls. Dark green represents 0% deuterium uptake (i.e., equal to the 0 % $D_2O$ control) and red represents maximally labeled (i.e., equal to the 100 % $D_2O$ control). The most dynamic regions are the insertion regions as these regions are maximally labeled after 1000 s. In contrast, the α/β hydrolase fold is more stable as witnessed by lower deuterium uptake.

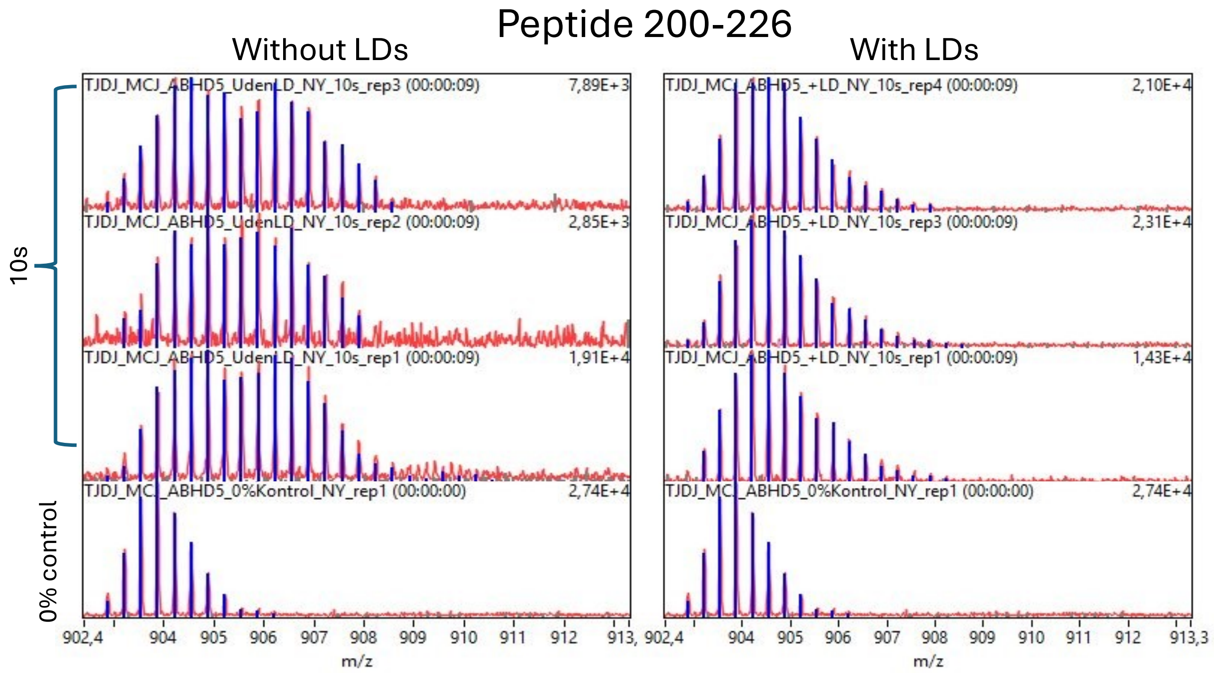


**Figure S4:** The conformational dynamics of the lid helix of ABHD5 in the presence and absence of LDs as measured by H/D exchange. ESI-MS spectra displaying the isotopic envelope of peptide 200-226 obtained from pepsin digestion of ABHD5 deuterated for 10 s at pH 7.3, 22°C in the absence or presence of LDs shown in left and right panel, respectively. The isotopic envelopes from triplicate experiments are shown, as well as the 0% control.

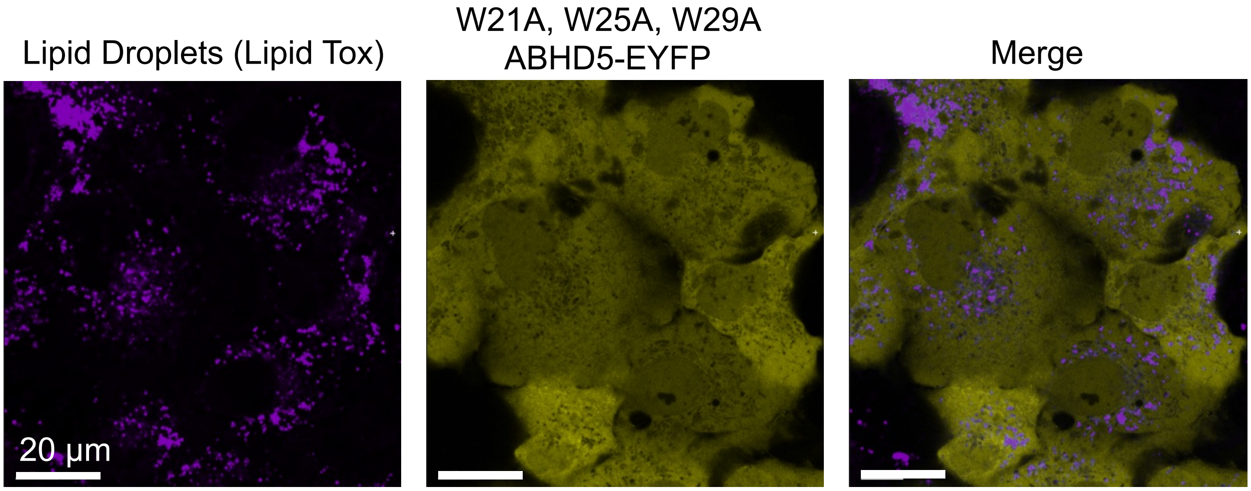


**Figure S5:** Mutation of W21, W25, and W29 prevents LD targeting of ABHD5 in Cos7 cells. (A) LDs were identified with LipidTox, a fluorescent neutral lipid stain. (B) W21A, W25A, W29A ABHD5 tagged with EYFP fails to colocalize to LD marked with LipidTox (C).

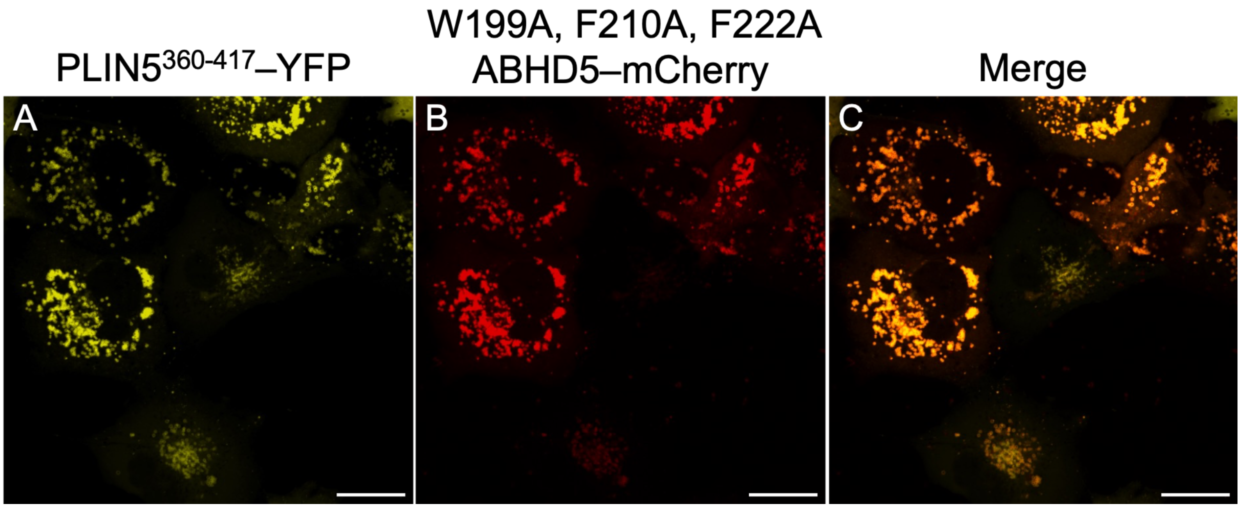


**Figure S6:** Confocal fluorescence imaging revealed that the triple mutant W199A, F210A, F222A ABHD5 was well expressed and properly folded by the Cos7 cells. This is demonstrated by the LD-targeting of the triple mutant ABHD5 when co-expressed with PLIN5 peptide that phenocopies the full length PLIN5 in regulating ABHD5 trafficking and lipolysis activation. (A) PLIN5360-417 concentrates on the LDs within live cells and (B) recruits the triple mutant to the LDs, (C) as shown colocalized in the color merged imaged. The scale bars represent 20 μm.

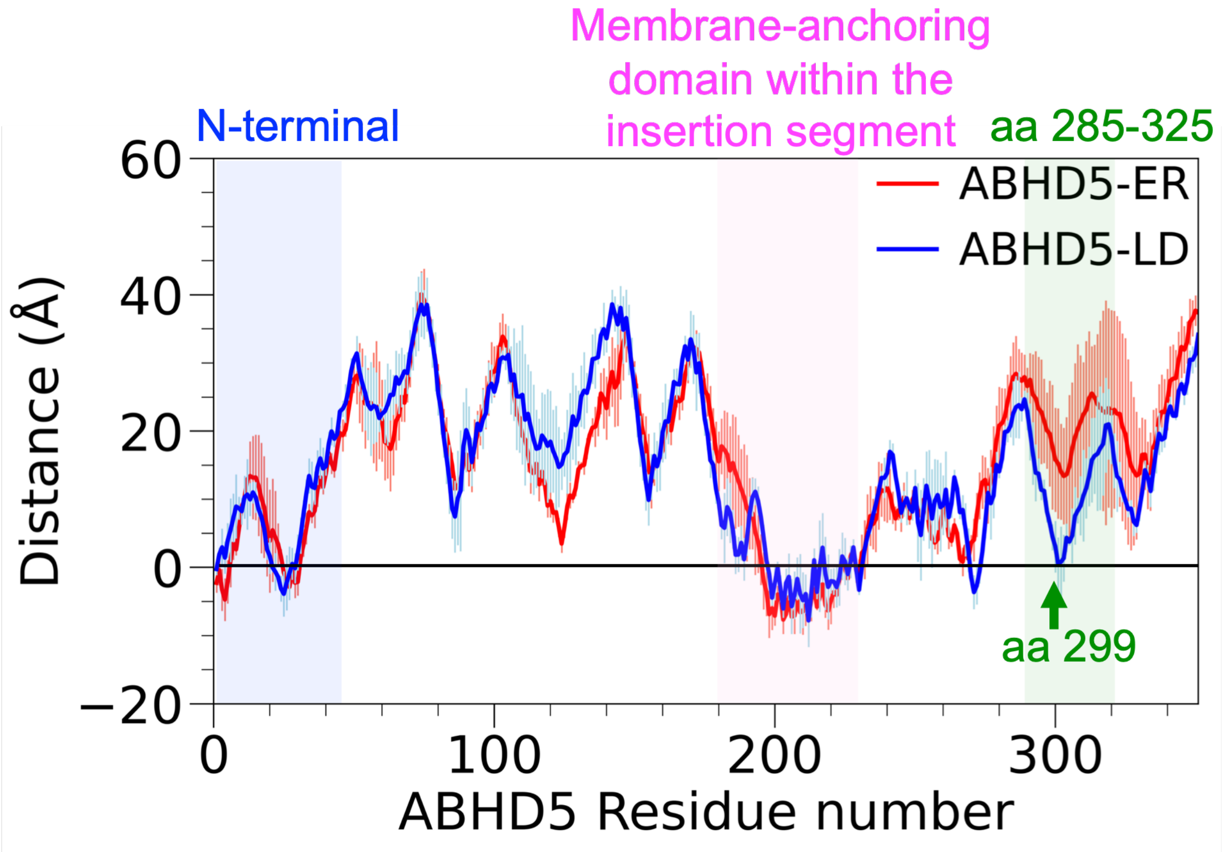


**Figure S7:** Distance between ABHD5 residues and the membrane surface. The green arrow indicates R299.

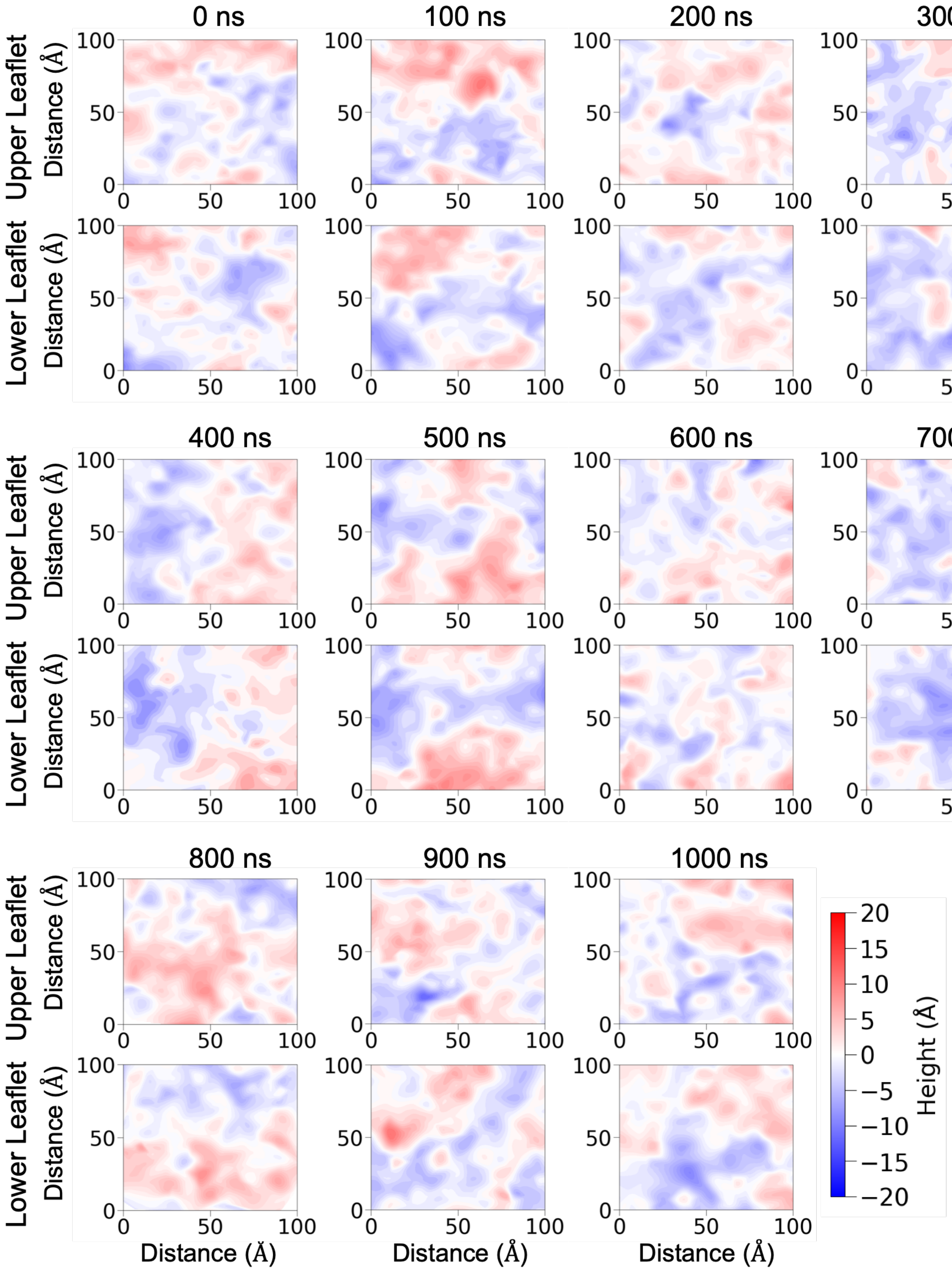


**Figure S8:** Changes in membrane surface height are shown at 100 ns intervals for the unbound ER membrane.

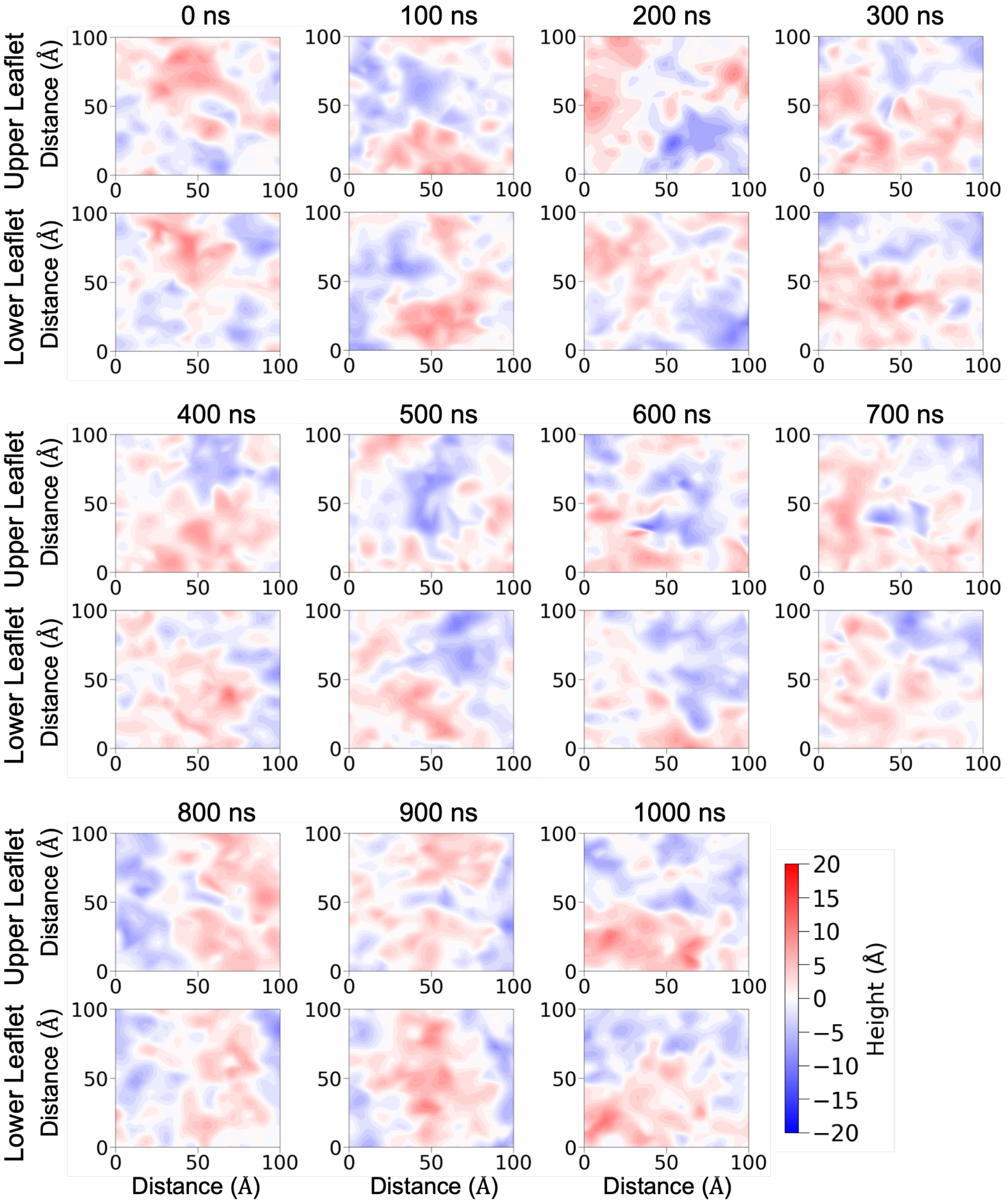


**Figure S9:** Changes in membrane surface height are shown at 100 ns intervals for the ABHD5-bound ER membrane.

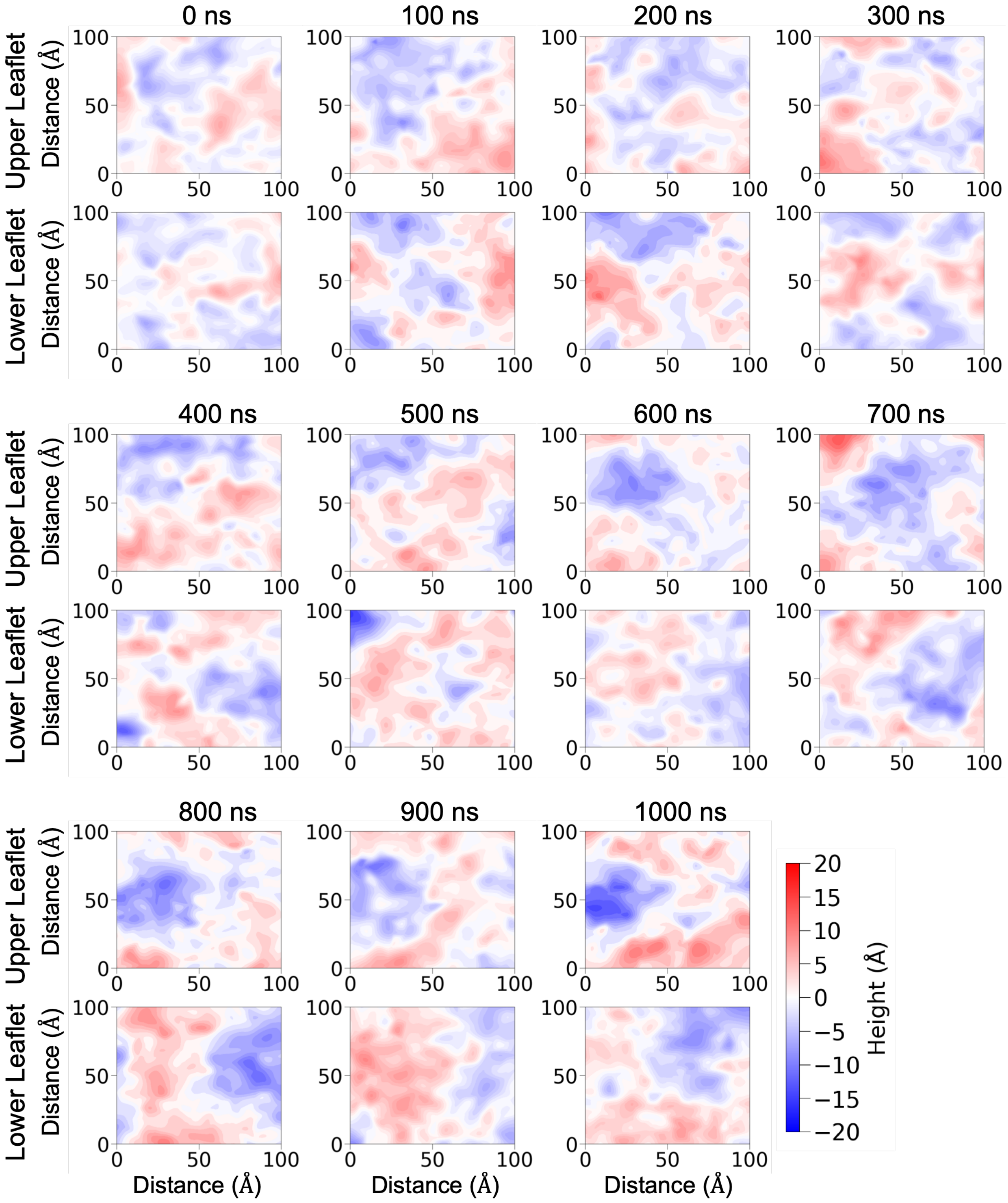


**Figure S10:** Changes in membrane surface height are shown at 100 ns intervals for the unbound LD membrane.

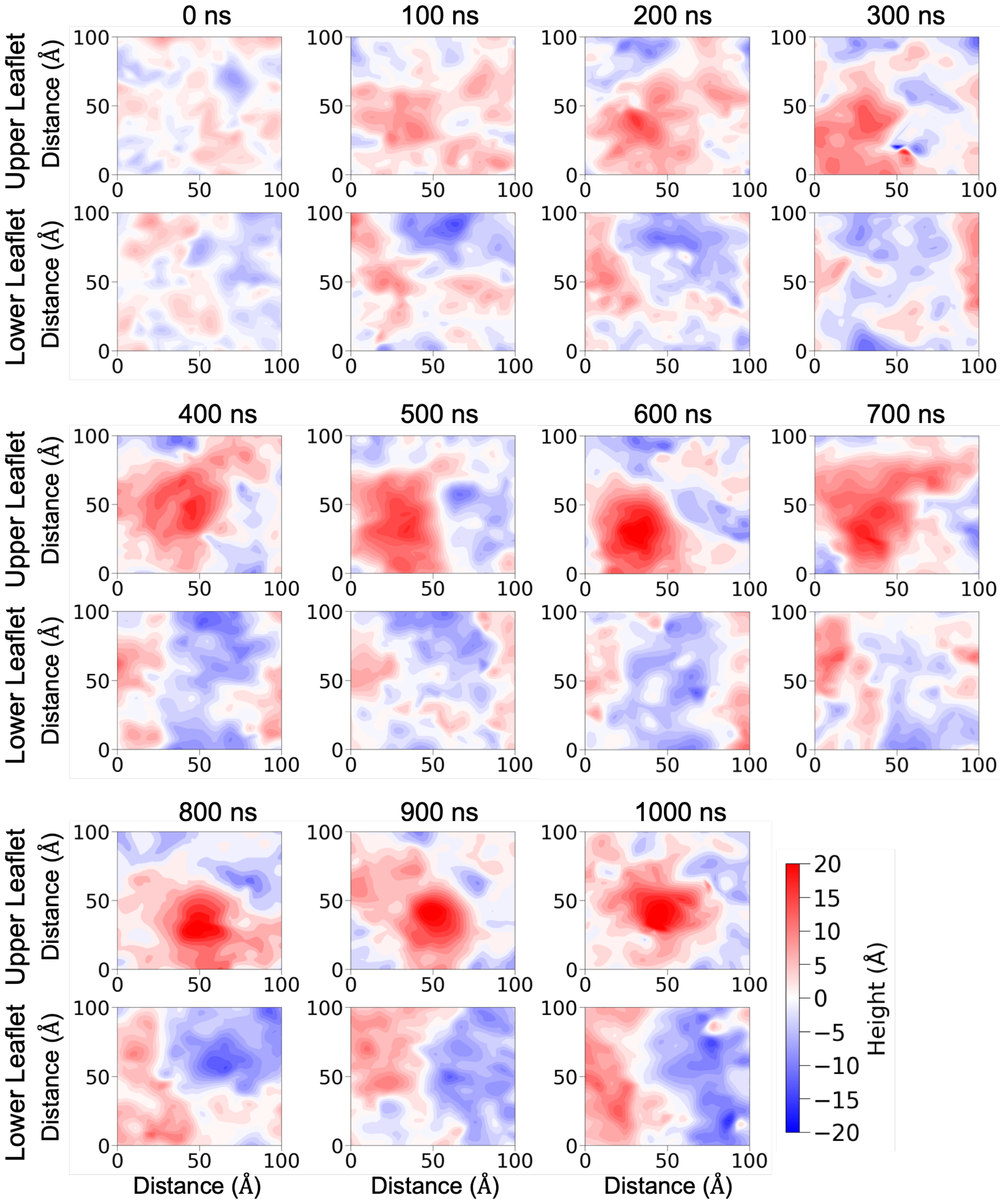


**Figure S11:** Changes in membrane surface height are shown at 100 ns intervals for the ABHD5-bound LD membrane.

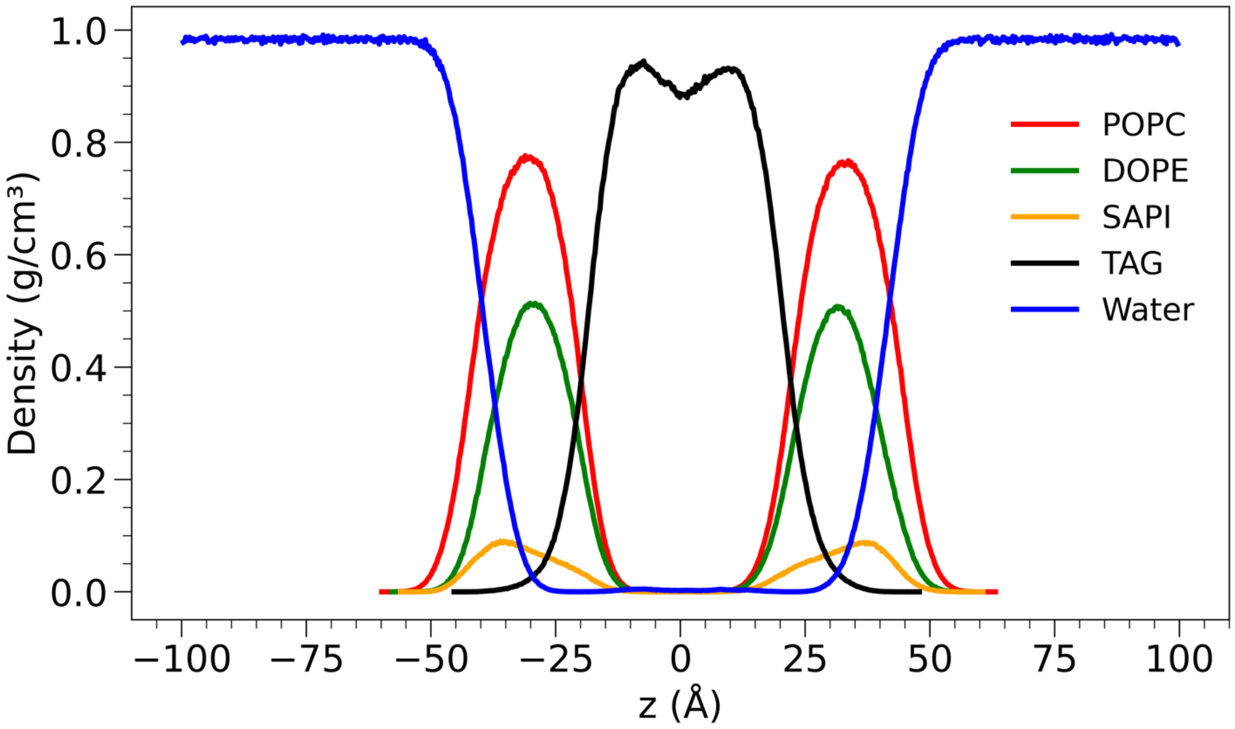


**Figure S12:** The mass density of each lipid in the LD membrane, where POPC, DOPE, SAPI, TAG, and water are represented in red, green, orange, black, and blue, respectively.

### References

1. Z. Zhang, D. L. Smith, Determination of amide hydrogen exchange by mass spectrometry: A new tool for protein structure elucidation. *Protein Sci.* **2**, 522–531 (1993).
